# Cell type deconvolution of bulk blood RNA-Seq to reveal biological insights of neuropsychiatric disorders

**DOI:** 10.1101/2023.05.24.542156

**Authors:** Toni Boltz, Tommer Schwarz, Merel Bot, Kangcheng Hou, Christa Caggiano, Sandra Lapinska, Chenda Duan, Marco P. Boks, Rene S. Kahn, Noah Zaitlen, Bogdan Pasaniuc, Roel Ophoff

## Abstract

Genome-wide association studies (GWAS) have uncovered susceptibility loci associated with psychiatric disorders like bipolar disorder (BP) and schizophrenia (SCZ). However, most of these loci are in non-coding regions of the genome with unknown causal mechanisms of the link between genetic variation and disease risk. Expression quantitative trait loci (eQTL) analysis of bulk tissue is a common approach to decipher underlying mechanisms, though this can obscure cell-type specific signals thus masking trait-relevant mechanisms. While single-cell sequencing can be prohibitively expensive in large cohorts, computationally inferred cell type proportions and cell type gene expression estimates have the potential to overcome these problems and advance mechanistic studies. Using bulk RNA-Seq from 1,730 samples derived from whole blood in a cohort ascertained for individuals with BP and SCZ this study estimated cell type proportions and their relation with disease status and medication. We found between 2,875 and 4,629 eGenes for each cell type, including 1,211 eGenes that are not found using bulk expression alone. We performed a colocalization test between cell type eQTLs and various traits and identified hundreds of associations between cell type eQTLs and GWAS loci that are not detected in bulk eQTLs. Finally, we investigated the effects of lithium use on cell type expression regulation and found examples of genes that are differentially regulated dependent on lithium use. Our study suggests that computational methods can be applied to large bulk RNA-Seq datasets of non-brain tissue to identify disease-relevant, cell type specific biology of psychiatric disorders and psychiatric medication.

## INTRODUCTION

One limitation of standard eQTL studies is that they generally use expression estimates from bulk tissue.^1, 2^ While this is informative, it has been shown that there are many cell type specific mechanisms driving biology,^3, 4^ which can be missed when looking at a collection of many cell types. In recent years, single cell RNA-Seq has allowed for the profiling of the gene expression of an individual cell, giving us a clearer picture of cell type gene expression. However, single cell RNA-Seq experiments are considerably more expensive than bulk RNA-Seq^5^. To leverage the advantages of each of these approaches, we can use methods to estimate cell type gene expression from bulk RNA-Seq expression.

There exist many methods^6, 7^ to estimate cell type expression from bulk RNA-Seq. Here, we elected to use CIBERSORTx^8^ and bMIND^9^ to estimate cell type proportions and cell type expression, respectively. Computational methods for analyzing bulk gene expression data have the potential for being advantageous in some applications as it is possible to obtain much larger sample sizes using bulk RNA-Seq instead of single cell RNA-Seq. While most single cell RNA-Seq studies have sample sizes in the range of several hundreds of cells from a small number of individuals, leveraging low-coverage bulk RNA-Seq allows us to obtain samples from hundreds to thousands of subjects.^10^ We used the low-coverage RNA-seq dataset described in Schwarz, et. al.^10^ as the primary dataset for analysis of cell type deconvolution in this study.

Associations between immune-related traits and neuropsychiatric disorders have been previously reported^11^, and we hypothesized that using blood-based expression can provide relevant information regarding the biology of such disorders.^12–14^ In this work we used cell type deconvolution methods to derive cell type specific estimates for gene expression from bulk blood RNA-seq, specifically within a cohort including psychiatric patients and controls of European ancestry. We used these results to conduct cell type cis-eQTL analyses, and compared the shared and unique cell type associations. We show that these cell type eQTL results derived from deconvoluted bulk RNA-Seq are consistent with eQTLs from scRNA-Seq. We performed colocalization analysis to find loci driving GWAS associations in either neuropsychiatric or blood-based traits and cell type gene expression. We go on to identify several examples of “opposite-effect” eQTLs, where a cell type eQTL signal demonstrates gene expression regulation in the opposite direction from that observed in a bulk eQTL study. Finally, we explored the effects of lithium use^15^ on cell type expression, and identified several cases of lithium-SNP interaction dictating presence of an eQTL.

## RESULTS

### Section 1: Computationally-derived cell type estimates are reliable

**Figure 1** provides a graphical abstract of the pipeline used in this study to generate putative cell-type specific eQTLs. To estimate cell type gene expression in whole blood, we analyzed bulk blood RNA-seq of bulk RNA-Seq (N = 1,730) using computational deconvolution tools. First, we estimated cell type proportions using the LM22 signature matrix and CIBERSORTx (**Figure 2A**). We found that these proportion estimates are consistent with standard white blood cell reference ranges,^16^ for which generally neutrophils have the highest abundance, lymphocytes (including T cells, B cells, natural killer (NK) cells combined) the second highest abundance, and monocytes the lowest abundance. However we note that blood cell type proportions vary across individuals depending on numerous factors such as medication use, current illness, and age.^17^ We confirmed that the proportions estimated via CIBERSORTx are consistent with the complete blood count measures taken in the clinic for a subset (N=143) of individuals in our dataset (**Supplementary Figure 1**). We observed a pearson correlation (R^2^) of 0.76 for cell type proportions estimated in neutrophils using CIBERSORTx and proportions measured in clinic, 0.85 for lymphocytes, and 0.48 for monocytes. These results suggest that the computationally estimated proportions are reliable.

**Figure 1:**
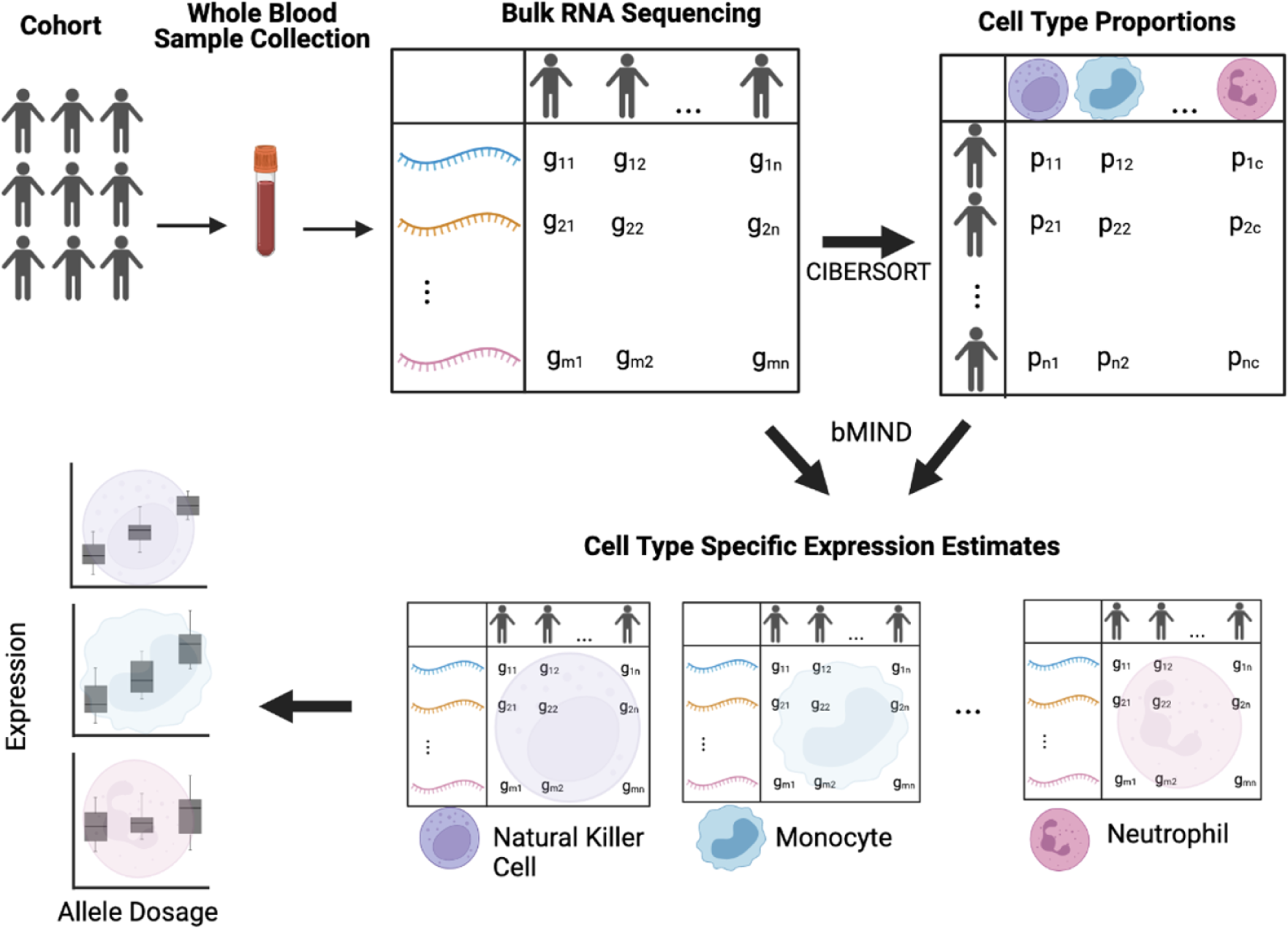
Graphical abstract of pipeline. Figure created in BioRender.

**Figure 2:**
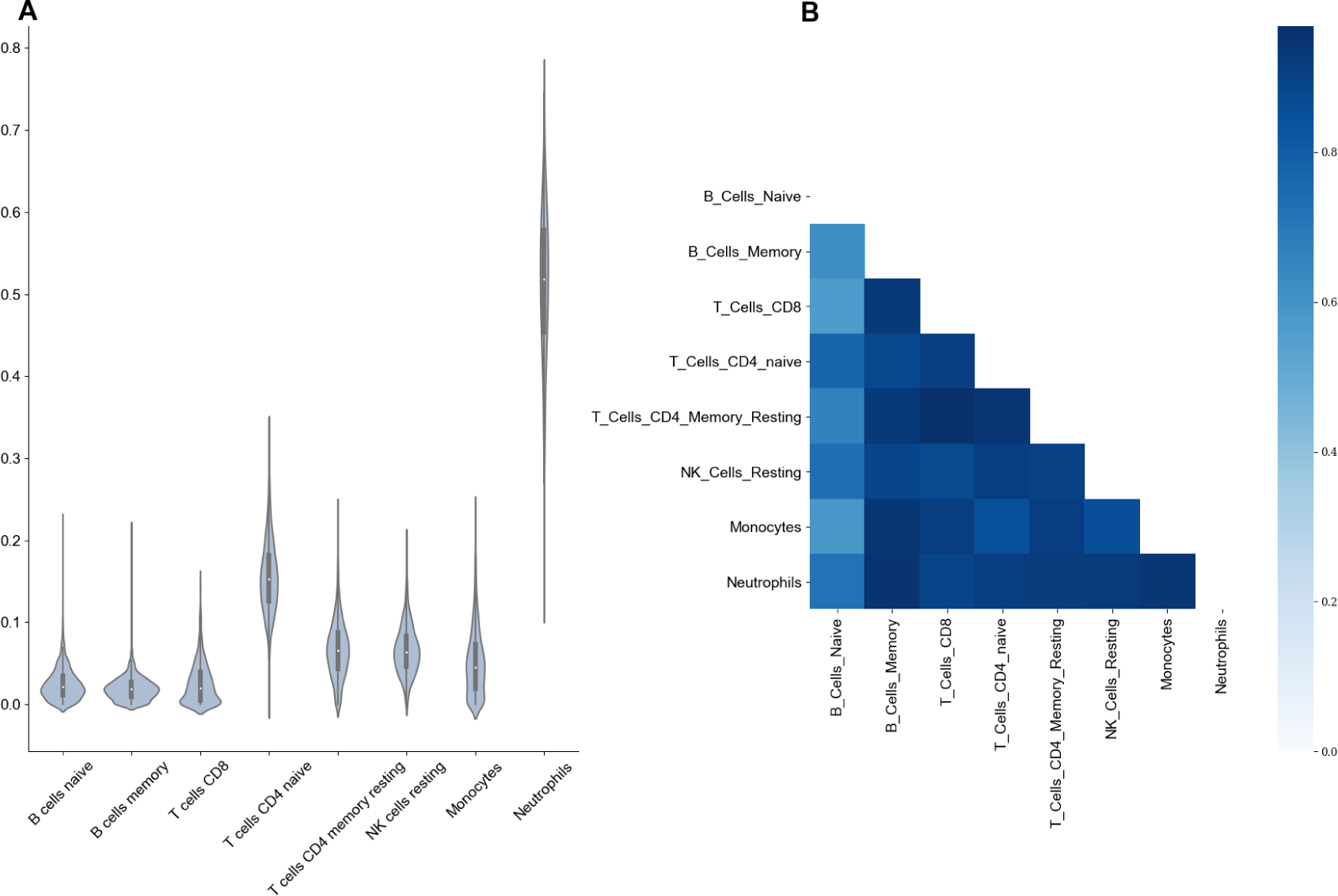
Cell type expression from computational deconvolution methods.

**Figure 2A:** Cell type proportion predictions from CIBERSORTx. - A violin plot showing the range of estimated cell type proportions for all 1730 individuals in each of the eight major cell types.

**Figure 2B:** R^2^ of expression between each cell type. - A heatmap of correlations (measured by R^2^ of mean expression across samples) between the eight main cell types captured by CIBERSORTx.

Next, we used these proportion estimates and bMIND expression deconvolution (**Methods**) to estimate cell type expression. Consistent with biological expectations, we found that correlation of estimated expression between different cell types is high, as all cell types are derived from the same tissue (**Figure 2B**). Next, we investigated whether computationally estimated cell type expression could successfully detect differences in expression between different cell types, despite there being a high correlation structure between different cell types. Principal component analysis confirmed that the major sources of variation in the dataset are attributable to differences in cell type expression (**Supplementary Figure 2**). These results suggest that using large cohorts of bulk RNA-Seq in blood, paired with computational deconvolution tools, can successfully detect differences in expression dependent on cell type composition.

Finally, we contrasted computationally-derived cell-type estimates with single cell RNA-Seq (scRNA-Seq) data.^18, 19^ We compared median TPM (transcripts per million) estimates across six cell types and find moderate correlation between the reference single-cell expression and computationally derived expression, ranging from R^2^ of 0.11 in naive B cells to R^2^ of 0.27 in CD8 T cells (**Supplementary Table 1** and **Supplementary Figure 3**). To further check how well computationally estimated expression compares to expression derived from scRNA-Seq, we correlated expression estimates between the two reference scRNA-Seq datasets in monocytes, the one cell type with data available in both reference datasets. We found that the median TPM of the 2,836 eGenes (genes with an associated eQTL) in both datasets have an R^2^ of 0.22, comparable to the R^2^ observed when comparing computationally estimated expression with scRNA-Seq.

### Section 2: Cell type eQTL analysis reveals more refined biological signal compared to bulk eQTL

Next, we performed eQTL analyses on the resulting cell type expression estimates to find evidence of genetic regulation of cell type expression. We restricted to the eight cell types with average proportion > 2% including: naive B Cells, memory B Cells, CD4 naive T Cells, CD4 memory T cells, natural killer cells, monocytes, and neutrophils. We conducted local-eQTL mapping with a 1 Mb window using QTLtools (**Methods**), to identify between 2,875 and 4,629 eQTL-genes (eGenes) with a significant association at FDR correction level of 5%, across the eight different cell types (**Figure 3A**). In total, we identified 5,752 eGenes with a significant association in at least one of the eight main cell types. We show that there exists a range of concordance of effect sizes for eGenes found in both the individual cell type analyses and the bulk eQTL analysis (**Figure 3B** and **3C**). This confirms findings from previous studies showing a strong shared genetic effect on gene expression across cell types. We observed that most eGenes are detected as significant in either just one, or all eight cell types (**Supplementary Figure 4**).

**Figure 3:**
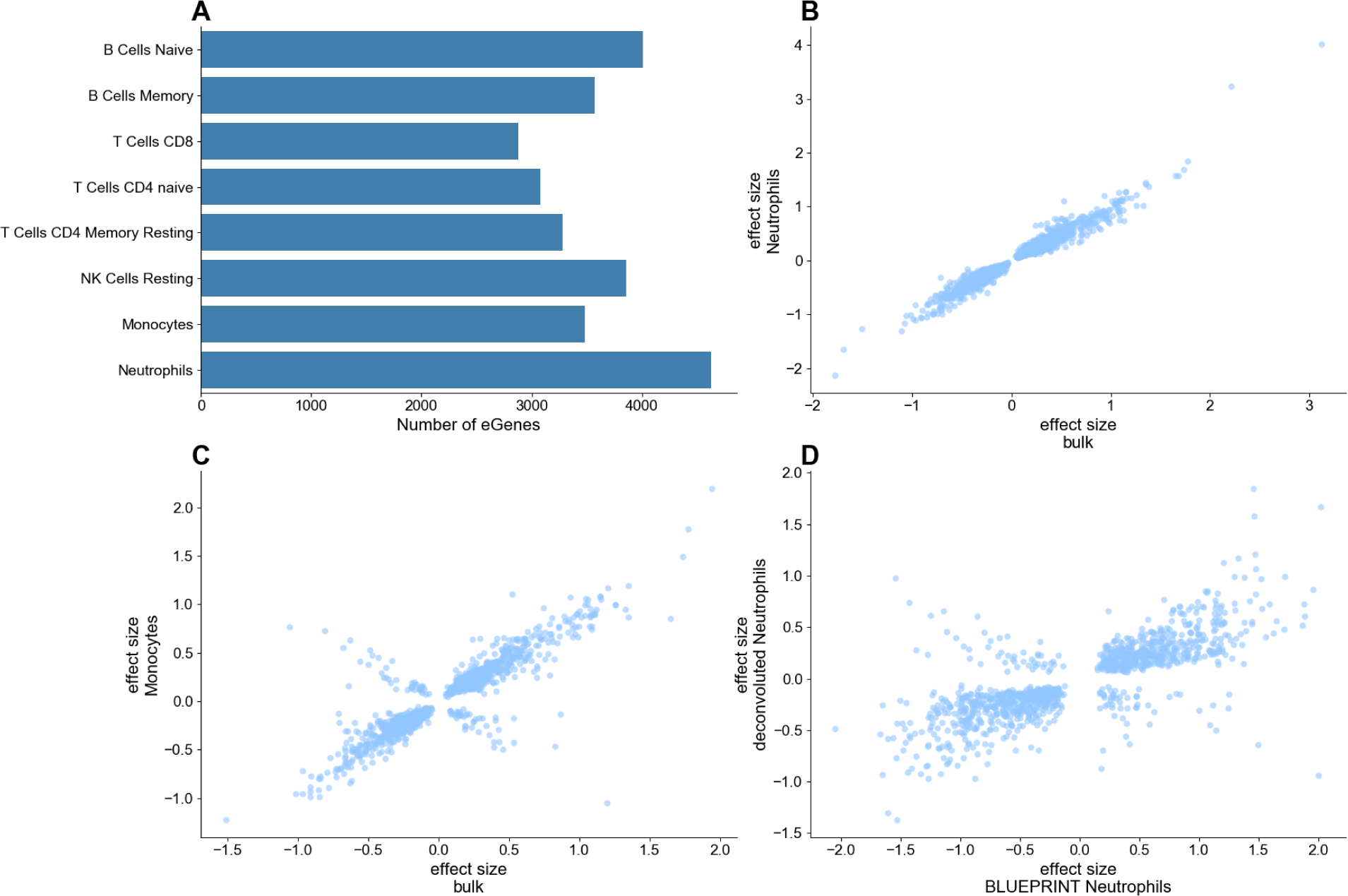
eQTLs per cell type, effect size correlation with reference dataset and bulk dataset.

**Figure 3A:** Number of associations identified per cell type. - Number of eGenes with a significant association identified for the eight major cell types detected by CIBERSORTx, using a FDR cutoff of 0.05.

**Figure 3B:** Comparison of effect size between shared cis-associations with Neutrophils. - Restricting to the eGenes with a significant association in both the bulk eQTL analysis and neutrophil eQTL analysis, we compare the estimated effect sizes of the most significant eQTL associations.

**Figure 3C:** Comparison of effect size between shared cis-associations with Monocytes. - Restricting to the eGenes with a significant association in both the bulk eQTL analysis and monocyte eQTL analysis, we compare the estimated effect sizes of the most significant eQTL associations.

**Figure 3D:** Comparison of effect sizes between shared cis-associations using reference single cell RNA-seq. Restricting to the eGenes with a significant association in both the BLUEPRINT reference neutrophil eQTL analysis and our neutrophil eQTL analysis, we compare the estimated effect sizes of the most significant eQTL associations.

Additionally, we found evidence of cell type “opposite-effect” eQTLs, where a SNP in a given cell type shows an association with the same eGene as detected using bulk RNA-Seq, but in the opposite direction. One such example is the eQTL for *FCGR3B* (Fc fragment of IgG receptor IIIb); while the bulk eQTL had an effect size of -1.3, the effect size in neutrophils and T cell types ranged between 0.49 and 0.86. Similarly, the eQTL for *MACF1* (Microtubule actin crosslinking factor 1) had effect sizes between -1.1 and -0.15 for the T cell types, versus effect sizes ranging between 0.21 and 0.28 for the bulk and remaining immune cell types. *MACF1* is known to be involved in neurite growth during brain development and has previously been linked to schizophrenia.^20^ These examples are especially interesting as it supports the idea that gene expression at the cell type level can uncover nuances of biological mechanisms that go undetected when only using bulk-level analyses. Similar effects have been observed in other studies using both single cell RNA-Seq^21^ and deconvoluted bulk RNA-Seq.^22^

To further validate these cell type eQTLs, we compared the results of this analysis with results from eQTL analysis using single cell RNA-Seq from the eQTLCatalogue and BLUEPRINT consortiums.^19, 23, 24^ We restricted to the protein coding genes identified as eGenes using the computational deconvolution approach. Generally, we found that the two approaches to cell type eQTL mapping show strong concordance. For example, in neutrophils, we found that 2,921 out of the 4,629 genes (63%) with a significant association using the computational deconvolution approach also had a significant association in using single-cell RNA-Seq, correcting at an FDR level of 5%. Among these eGenes, comparing the association with the same leading SNP in both of these datasets (**Figure 3D**), we observed a correlation (R^2^) of 0.66 between their effect sizes. Similar effect size correlations, for T cells CD4, B cells, and monocytes are shown in **Supplementary Figure 5**. This suggests that the computational deconvolution approach to large scale bulk RNA-Seq projects can be used to obtain accurate cell type eQTL estimates.

### Section 3: Integration of cell type specific eQTL with brain and blood trait GWAS

For every gene with a significant eQTL, we used FUSION^25^ to estimate the gene expression heritability across each of the contexts, or the proportion of variance in gene expression explained by variance in genetics. Only those genes with significant heritability after five-fold cross validation per each context were retained for further analysis. **Table 2** provides the summarized statistics of the significantly heritable genes and the gene with highest estimated SNP-heritability per cell type. An advantage of investigating eQTLs at the cell type level is that it provides a more precise view of biological mechanisms driving the association between gene expression and phenotype. In order to investigate whether there exists variants that drive both the expression of genes in a specific cell type and a GWAS trait, we conducted Transcriptome Wide Association Study (TWAS)^25^ and colocalization^26^ analyses using the significant ct-eQTLs from the eight main cell types previously mentioned, and GWAS of several neuropsychiatric and blood-based phenotypes. **Figure 4A** provides an overview of the overlap across the contexts, both for brain-related and blood-based traits.

**Table 1:**
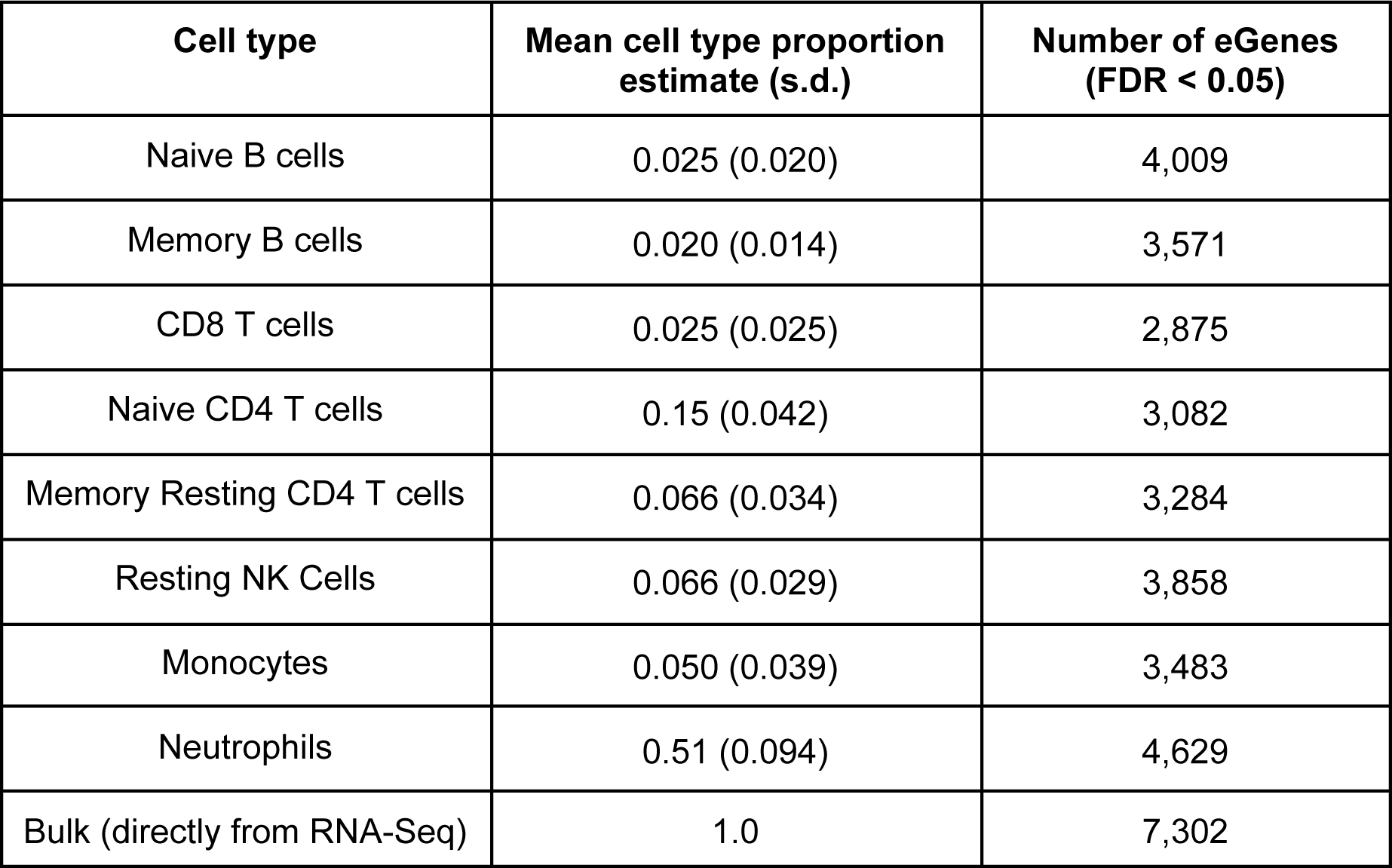
Cell type proportion estimates from CIBERSORTx and number of eQTLs per cell type.

| Cell type | Mean cell type proportion estimate (s.d.) | Number of eGenes (FDR < 0.05) |
| --- | --- | --- |
| Naive B cells | 0.025 (0.020) | 4,009 |
| Memory B cells | 0.020 (0.014) | 3,571 |
| CD8 T cells | 0.025 (0.025) | 2,875 |
| Naive CD4 T cells | 0.15 (0.042) | 3,082 |
| Memory Resting CD4 T cells | 0.066 (0.034) | 3,284 |
| Resting NK Cells | 0.066 (0.029) | 3,858 |
| Monocytes | 0.050 (0.039) | 3,483 |
| Neutrophils | 0.51 (0.094) | 4,629 |
| Bulk (directly from RNA-Seq) | 1.0 | 7,302 |

**Table 2:** FUSION heritability results. Number of Sig. Genes refers to the number of genes that remain significantly (P<0.05) heritable after five-fold cross validation. Q1 = first interquartile, Q3 = third interquartile. Overall, the bulk data shows higher heritability estimates across each of the statistics. Of note is that every gene listed is distinct for each context, including genes that are relevant to neuronal function, such as *NSG1* (neuronal vesicle trafficking associated), *CAMKK2* (calcium dependent kinase, involved in neuronal differentiation and synapse formation) and *BTG1* (B-cell translocation gene 1, found to be involved in neural stem cell renewal).^61^

| Panel | Number of Sig. Genes | Min | Q1 | Median | Mean | Q3 | Max | Gene with Max h2 |
| --- | --- | --- | --- | --- | --- | --- | --- | --- |
| **Bulk** | 5,113 | 0.0041 | 0.026 | 0.055 | 0.096 | 0.12 | 0.728 | *TRBV28* |
| **B Cells Memory** | 2,541 | 0.0035 | 0.024 | 0.044 | 0.075 | 0.093 | 0.68 | *BTG1* |
| **B Cells Naive** | 1,552 | 0.0056 | 0.024 | 0.045 | 0.078 | 0.095 | 0.579 | *PI16* |
| **Monocytes** | 2,431 | 0.0052 | 0.025 | 0.045 | 0.077 | 0.095 | 0.584 | *NSG1* |
| **NK Cells Resting** | 2,763 | 0.0042 | 0.024 | 0.045 | 0.078 | 0.098 | 0.61 | *BCAT1* |
| **Neutrophils** | 1,605 | 0.0056 | 0.025 | 0.048 | 0.083 | 0.10 | 0.69 | *CAMKK2* |
| **T Cells CD4**<br>**Memory Resting** | 1,989 | 0.0057 | 0.026 | 0.047 | 0.080 | 0.099 | 0.63 | *SBF2* |
| **T Cells CD8** | 2,033 | 0.0057 | 0.024 | 0.042 | 0.069 | 0.081 | 0.63 | *FGFBP2* |
| **T Cells CD4 Naive** | 2,147 | 0.0053 | 0.024 | 0.044 | 0.075 | 0.092 | 0.56 | *CROT* |

**Figure 4:**
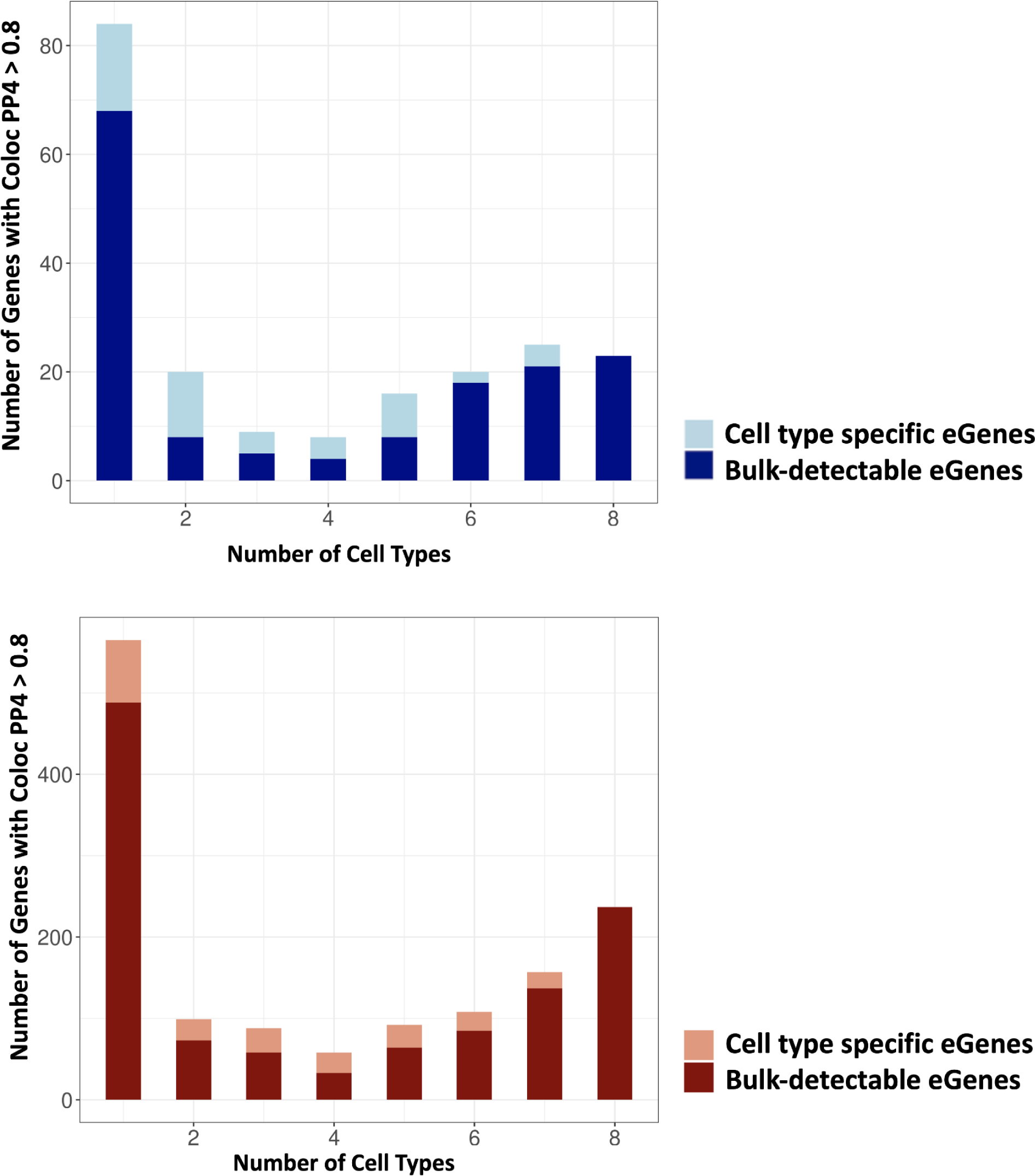
Colocalization and enrichment analyses of cell type specific eQTLs.

**Figure 4A:** (top) Number of genes with coloc PP4>0.8 across contexts in neuropsychiatric traits. (bottom) Number of genes with coloc PP4>0.8 across contexts in blood-based traits.

**Figure 4B:**
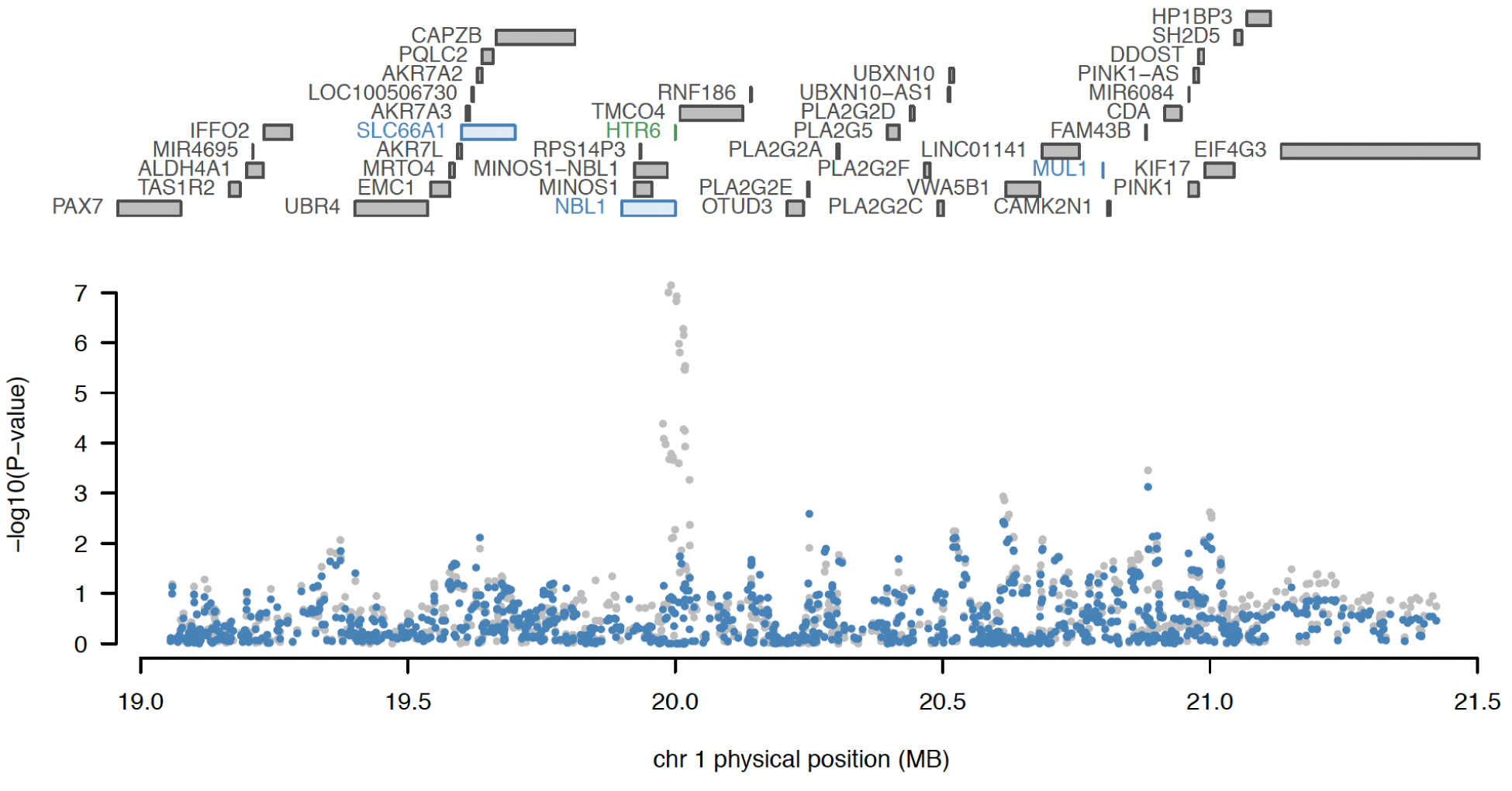
Conditional analysis of *HTR6* expression in memory B cells. All genes in the locus are included in the top panel, with marginally TWAS associated genes highlighted in blue, and those jointly significant (*HTR6*) in green. The bottom panel includes a Manhattan plot of the GWAS data before (gray) and after (blue) conditioning on the imputed expression of *HTR6* in memory B cells. Figure generated by FUSION.post_process.R script.

GWAS for neuropsychiatric traits tested include: BP,^27^ SCZ,^28^ major depressive disorder (MDD),^29^ alcohol dependence,^30^ cannabis use disorder,^31^ migraines,^32^ insomnia,^33^ attention-deficit/hyperactivity disorder (ADHD),^34^ and Alzheimer’s disease.^35^ In total there were 710 eGenes found to be associated only in bulk and no other cell type, and 168 eGenes found to be associated in one or more cell types and not in the bulk (**Table 3**). Regarding colocalization, in total there were 68 eGenes found to have colocalized SNPs between expression and trait only in the bulk and no other cell type, and 50 eGenes found only in one or more cell types and not in the bulk (**Table 3**).

**Table 3:** TWAS & Colocalization neuropsychiatric trait results. Shared refers to the number of significant (FDR < 0.05) genes that are in common with the bulk TWAS-significant gene set, whereas unique refers to those that are not present in the bulk TWAS-significant gene set.

| **Cell Type** | **Significant**<br>**eGenes** | **Num Significant**<br>**TWAS Genes,**<br>**Shared** | **Num Significant**<br>**TWAS Genes,**<br>**Unique** | **Num Genes with**<br>**Coloc PP4>0.8,**<br>**Shared** | **Num Genes with**<br>**Coloc PP4>0.8,**<br>**Unique** |
| --- | --- | --- | --- | --- | --- |
| **B Cells Naïve** | 4,009 | 90 | 43 | 43 | 13 |
| **B Cells Memory** | 3,571 | 142 | 58 | 62 | 25 |
| **T Cells CD8** | 2,875 | 108 | 50 | 50 | 15 |
| **T Cells CD4 Naïve** | 3,082 | 120 | 46 | 56 | 19 |
| **T Cells CD4 Memory**<br>**Resting** | 3,082 | 115 | 43 | 55 | 22 |
| **NK Cells Resting** | 3,858 | 156 | 72 | 73 | 21 |
| **Monocytes** | 3,483 | 126 | 52 | 62 | 24 |
| **Neutrophils** | 4,629 | 76 | 35 | 35 | 9 |
| **Bulk** | 7,302 | 906 | / | 155 | / |

Of the 50 eGenes found to have a colocalization posterior probability with the same variant impacting both gene expression and the GWAS trait (PP4>0.8) in a cell type but not in the bulk, half have a higher median TPM across the GTEx v8 brain tissue types than in GTEx whole blood. This suggests that these genes are relevant for brain functions despite being detected in immune cell type specific expression estimates. An example of one such gene is *HTR6*, a serotonin receptor targeted by certain antidepressant and antipsychotic medication, found to be strongly associated and colocalized with BP in the most recent Psychiatric Genomics Consortium (PGC) study on bipolar disorder^27^ which used brain-derived gene expression weights from the PsychENCODE project.^36^ Conditioning on *HTR6* memory B cell-specific expression using FUSION completely removed the significant GWAS signal at this locus, suggesting that the genetic factor driving gene expression also encompasses the BP association signal (**Figure 4B**). The same held true for other immune cell types in which *HTR6* was colocalized with BP, including naive B cells and CD4 T cells. This demonstrates the utility of using cell type deconvolution methods in large cohorts of an easily-accessible tissue like blood, since it is able to capture gene expression regulation relevant in brain cell types that otherwise are not detectable in bulk blood eQTLs.

GWAS for blood-based traits tested include: systemic lupus erythematosus^37^ (an autoimmune disorder), mean corpuscular volume, mean corpuscular hemoglobin,^38^ red blood cell width distribution, monocyte count, eosinophil count, lymphocyte count, platelet count, white blood cell count, and red blood cell count.^39^ In total there were 1,765 eGenes found to have associations only in bulk and no other cell type, and 493 eGenes found only in one or more cell types and not in the bulk (**Table 4**). Regarding colocalization, in total there were 488 eGenes found only in the bulk and no other cell type, and 229 eGenes found only in one or more cell types and not in the bulk (**Table 4**).

**Table 4:** TWAS & Colocalization blood-based trait results. Shared refers to the number of significant (FDR < 0.05) genes that are in common with the bulk TWAS-significant gene set, whereas unique refers to those that are not present in the bulk TWAS-significant gene set.

| **Cell Type** | **Significant**<br>**eGenes** | **Num Significant**<br>**TWAS Genes,**<br>**Shared** | **Num Significant**<br>**TWAS Genes,**<br>**Unique** | **Num Genes with**<br>**Coloc PP4>0.8,**<br>**Shared** | **Num Genes with**<br>**Coloc PP4>0.8,**<br>**Unique** |
| --- | --- | --- | --- | --- | --- |
| **B Cells Naïve** | 4,009 | 922 | 164 | 289 | 78 |
| **B Cells Memory** | 3,571 | 1,582 | 207 | 511 | 106 |
| **T Cells CD8** | 2,875 | 1,276 | 168 | 414 | 93 |
| **T Cells CD4 Naïve** | 3,082 | 1,349 | 183 | 445 | 88 |
| **T Cells CD4 Memory**<br>**Resting** | 3,082 | 1,257 | 150 | 419 | 80 |
| **NK Cells Resting** | 3,858 | 1,712 | 254 | 557 | 119 |
| **Monocytes** | 3,483 | 1,484 | 212 | 484 | 113 |
| **Neutrophils** | 4,629 | 969 | 159 | 331 | 60 |
| **Bulk** | 7,302 | 3,893 | / | 1,175 | / |

Within the blood-based traits we again found examples of opposite-sign effects in certain cell types when compared to the bulk. For example, when considering systemic lupus erythematosus (SLE) as a trait, we found for the *IRF5* gene, natural killer cells have a TWAS Z-score of -10.7 whereas the bulk has a score of +3.91, suggesting distinct mechanisms that are dependent on the cell type context. IRF5 (interferon regulatory factor 5) is known to be implicated in SLE,^40, 41^ though the exact mechanism by which it is dysregulated in the context of disease remains unknown.

See the TWAS Supplementary Tables to view all FUSION TWAS and colocalization results.

### Section 4: Lithium-dependent genetic regulation of gene expression

Given the large number of BP probands in our study sample, we were interested to see whether there were BP-specific effects that could be observed using cell type deconvoluted expression. Since lithium is the most commonly used drug to treat these patients and it has also been established that lithium use has an effect on the blood transcriptome,^42, 43^ we hypothesized that lithium-dependent genetic regulation of the blood transcriptome may exist. Among the 1,045 bipolar disorder patients in this cohort, 709 were taking lithium at the time of blood draw (“Lithium-User”) and 336 were not (“Lithium Non-User”).

When stratifying by cases versus controls (with all BP and SCZ individuals included as cases), we found significant differences in cell type proportion for CD4 T cells (p=1.8e-7, higher in controls), natural killer resting cells (p=1.2e-7, higher in controls), and neutrophils (p=2.3e-8, higher in cases). Next, considering only the cases of BP, we stratified those who use lithium versus those who do not, and found significant differences in cell type proportion for CD4 naive T cells (p=8e-4, higher in non-users), CD4 memory T cells (p=4e-4, higher in non-users), natural killer resting cells (p=3e-4, higher in non-users), and neutrophils (p=1.5e-9, higher in users). However, when we only include lithium non-users within the BP cases, and compare those against the controls, we found no significant differences in proportion for any of the cell types. See **Supplementary Figure 6** for example plots of all three tests using neutrophils. This suggests that the use of lithium within the BP cases drives these differences in cell type proportion, rather than disease status itself, consistent with previous findings.^43^

We validated the effect of lithium use on blood cell types in a separate cohort of individuals who had electronic health data from the University of California, Los Angeles ATLAS Community Health Initiative.^44, 45^ Specifically, we included self-reported European patients with a PheCode for bipolar disorder who also had laboratory test orders for complete blood counts and noted whether they had a prescription order for lithium (n=1302 with lithium, n=6208 without). In comparing the neutrophil count between BP patients who had never been prescribed lithium (or before they were prescribed lithium) and those who had a prescription order for lithium, we found that there was a significant (logistic regression p=2.09e-07) elevation of neutrophils in patients with a prescription for lithium (**Supplementary Figure 7**). Furthermore, for a subset of BP patients within the ATLAS dataset, we also have records for neutrophil counts both before and after the patient was prescribed lithium. Using a Wilcoxon-signed rank test with continuity correction, we found a significant difference between the neutrophil counts between the two groups (p=0.0228) when including individuals of any ancestry (n=376), though when restricting to only European individuals (n=229), the significant difference is lost (p=0.2) (**Supplementary Figure 7**). The replication of this finding in this large external dataset provides further evidence to suggest that cell type proportion is impacted by lithium usage, though the implications of this are yet to be understood.

Next, we investigated whether estimated cell type expression is a significant predictor for case/control status or lithium use. Restricting to the genes with the highest variance in each cell type, we built logistic regression models to separately predict case/control status and lithium use, including the same covariates as the previous proportion-based models. However, we find that gene expression does not provide additional predictive value over the cell type proportions for either case/control status or lithium use.

To investigate lithium-dependent genetic regulation, we performed an interaction model eQTL scan between lithium users and nonusers, testing whether there exist SNPs whose cell type or cell type specific expression regulation is dependent on the presence of lithium. To do this, we included an interaction term for the genotypes and lithium status in the regression model (**Methods**). Using bulk expression, we only identified one gene with such an association (FDR p-value < 0.10). With cell type expression derived from bMIND, we identified as many as 34 such eGenes (in monocytes), and a total of 110 examples of genes (Li-eGenes) that show differential regulation of cell type expression, compared to just one gene that shows differential regulation of bulk expression (**Supplementary Table 3**). We found that 97 of the eGenes that have significant differential lithium regulation exhibit opposite effect sizes between the lithium user and nonuser groups, at the cell type level. The remaining 13 Li-eGenes show same direction effect sizes between the lithium user and nonuser groups, with significantly different magnitudes. For example, in naïve B cells, *KITLG* (ENSG00000049130) shows opposite effect eQTLs based on rs11104703 (**Figure 5A**). While in monocytes we see that *TNFRSF11A* (ENSG00000105641) shows differential effect size, in the same direction, based on rs79143095 (**Figure 5B**). Due to the large number of samples used in this analysis, we are powered to detect small differences, like these.

**Figure 5:**
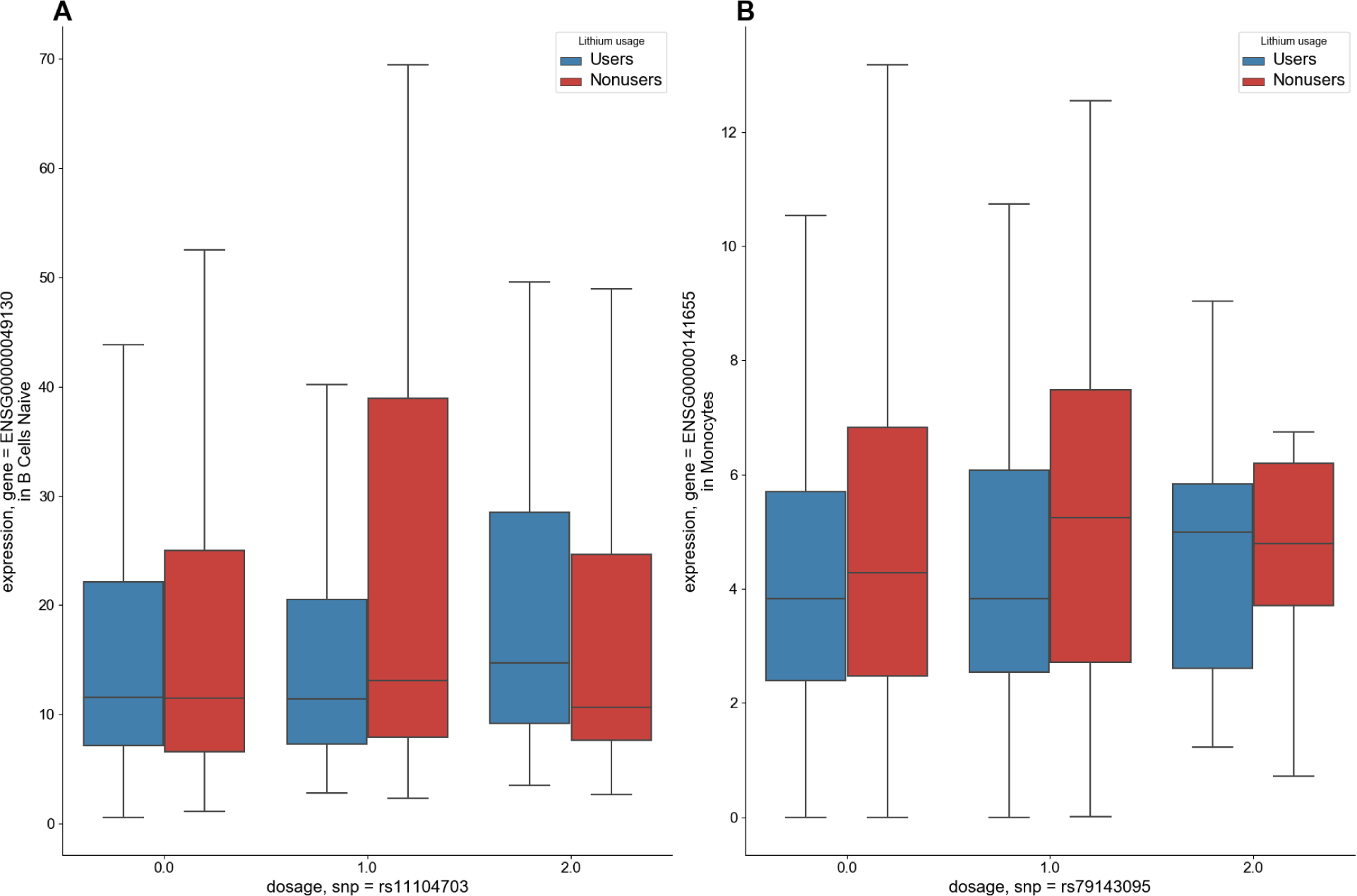
Lithium user vs non-user analyses.

**Figure 5A:** Boxplots showing the expression of *KITLG* (ENSG00000049130) in naïve B cells, stratified by dosage of SNP rs11104703 in lithium users versus nonusers. **5B:** Boxplots showing the expression of *TNFRSF11A* (ENSG00000105641) in monocytes, stratified by dosage of SNP rs79143095 in lithium users versus nonusers.

**Figure 5C:**
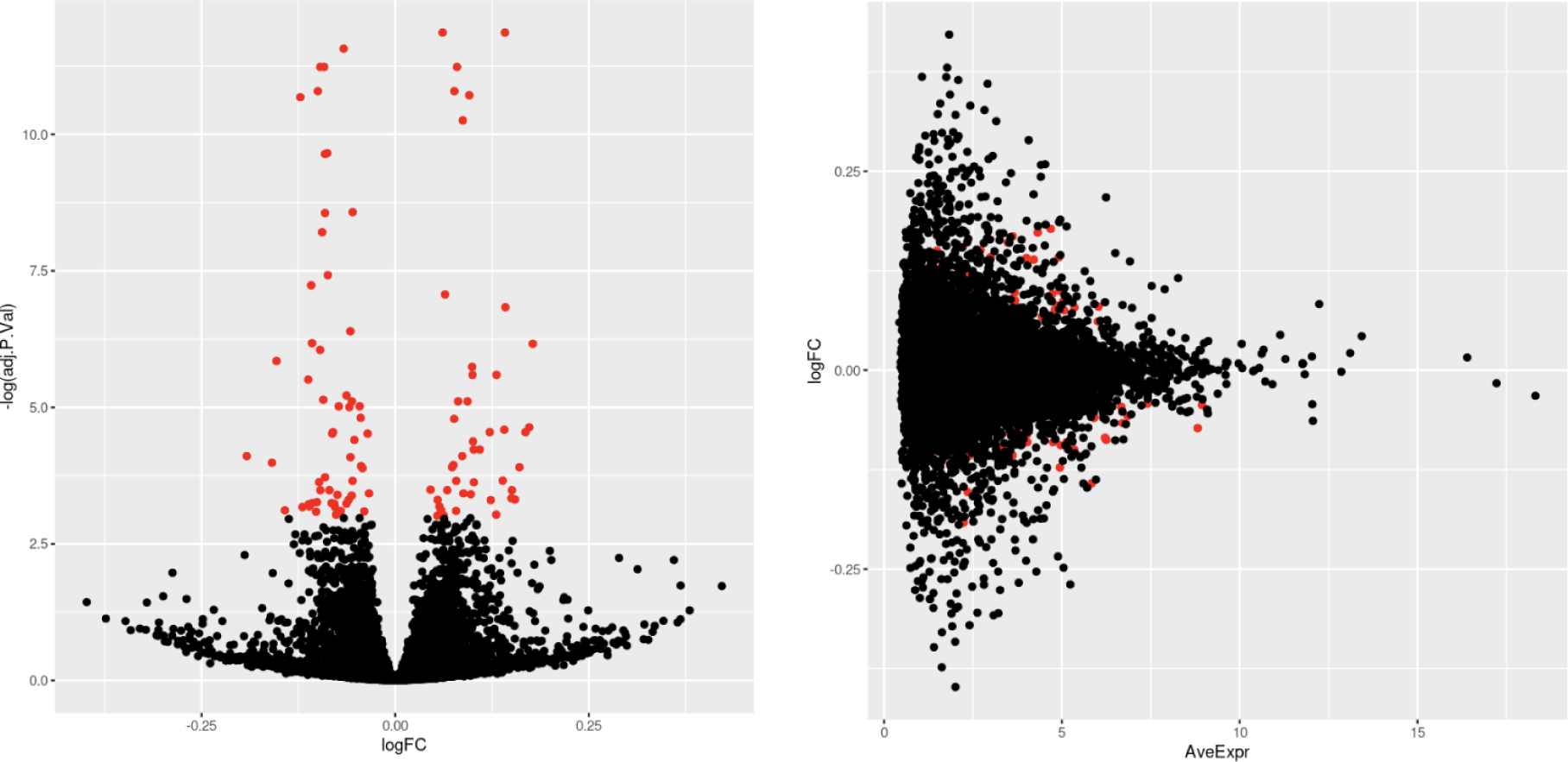
Differential gene expression results for lithium users vs lithium non-users: (left) Volcano plot which highlights differentially expressed genes (FDR < 0.05) in red (N=100 total differentially expressed genes). (right) Average expression of each gene vs the log fold change (logFC) of each gene, with differentially expressed genes highlighted in red.

In order to directly measure expression differences between lithium users and nonusers, we conducted a differential expression analysis test using limma^46^ initially in the bulk dataset (**Methods**). Comparing the two groups, we tested 17,194 genes from bulk expression measures. We found 100 genes with evidence of differential expression in bulk (FDR < 0.05), with log fold changes of the significant genes ranging from -0.191 to 0.177, suggesting low impact of lithium on differential expression (**Figure 5C**). Out of the 100 differentially expressed genes found here, 33 were previously reported in Krebs, et. al,^43^ a significant overlap according to Fisher’s exact test (OR = 6.43, p = 4.74e-14). Overlapping genes include *FBXL2* - a gene highly expressed in the brain and involved in neuronal signaling, and *CNTNAP3* - which mediates interactions between neurons and glial cells. See Differential Expression Supplementary Tables for full lithium differential expression results.

Though previous studies have not found substantial evidence of differential expression in the blood transcriptome between cases of BP or SCZ and controls,^43, 47^ we were interested in investigating this within our own cohort given the uniquely large sample size. Using the bulk RNA-seq and the same 17,194 genes selected in the lithium-user differential expression analysis, we found 64 genes with FDR < 0.05, of which nine genes overlapped with the significant genes found in the lithium analysis. Log fold changes of the significant genes ranged only from -0.126 to 0.104, suggesting that if these genes are truly a result of disease status, the differences are minimal (**Supplementary Figure 8**). See Differential Expression Supplementary Tables file for full case/control differential expression results.

For the cell-type specific differential expression analyses, we leveraged the differential expression function available through the bMIND software. In the case-control analysis, we found four differentially expressed genes in Neutrophils (FDR<0.05), including *TSPAN2* and *CFAP45* both of which were reported in the Krebs et. al. lithium differential expression study.^43^ We found 24 differentially expressed genes in memory B cells, and 21 in naive B cells (with 18 differentially expressed genes in common between the two B cell types). Interestingly, when conducting the lithium user versus non-user analysis, we did not find any differentially expressed genes in any cell type. While this may be a result of the smaller sample set used in the lithium analysis as compared to the case-control analysis, it also may reflect that the effects of lithium are only found at the bulk level due to its impact on cell type composition, rather than changes in gene expression within individual cell types. See Differential Expression Supplementary Tables file for q-values of all cell type specific differential expression results.

## DISCUSSION

We show that cell type deconvolution of bulk blood RNA-seq provides novel insights not only for immune-relevant biology, but also neuropsychiatric disease biology. While bulk eQTLs tend to provide a greater number of associations overall, we find that cell type specific eQTLs provide unique associations not otherwise detectable in bulk. Many of these unique cell type associations have high expression in brain tissue types, and harbor several example genes that have been previously implicated in BP TWAS^27^ studies using brain tissue. This demonstrates that large cohorts of an easily accessible tissue like blood is useful for deciphering biology for brain-related phenotypes when cell type deconvolution is applied. An important caveat, however, is that the associations with brain-related traits found in this study are most likely to be shared genetic mechanisms between blood cell types and brain cell types, rather than blood cell type-specific biology.

Considering the BP TWAS results alone, there were 82 total eGenes with an opposite direction of effect in a cell type than in the bulk eQTL analysis (defined as having an opposite-sign TWAS Z-score for the same gene and the same trait). For example, we found 63 eGenes, significantly associated with BP, that have an opposite direction of effect in CD8-T cells when compared to bulk expression. *ARID5A*, a gene implicated in the most recent PGC bipolar disorder TWAS^27^ is one example of these genes. In the bulk expression the TWAS Z-score of *ARID5A* and bipolar disorder is -4.99 (TWAS Z-score -5.32 in PGC BP study), whereas in CD8-T cells it is +6.02. This gene was also found to be colocalized with PP4>0.8 in the CD8 T Cell test, though it does not pass the colocalization threshold in the bulk test or PGC3 BP test. The same is true for *ARID5A* in CD4 memory resting T cells (TWAS Z-score +6.56). Similarly, the methyltransferase gene *WDR82* in CD4 Naive T cells has a positive (TWAS Z-score +3.72) association with BP, whereas the bulk expression has a negative (TWAS Z-score -3.98) association at the same locus (TWAS Z-score -6.75 in PGC BP study). There are many such examples of these genes across each of the cell types and the various traits that we examined.

Examples of novel BP-associated genes were also discovered, including *RILPL2*, found to be colocalized in the context of memory B cells, monocytes, natural killer resting cells, and CD8 T cells, but not in the bulk. This gene is highly expressed in whole blood in adults (median TPM 27.42 in GTEx), but is also crucial for dendritic-spine morphogenesis in developing neurons^48^ Similarly, *CAMKK2* (calcium/calmodulin dependent protein kinase kinase 2), a gene found to be colocalized in the context of monocytes, neutrophils, and CD4 T cells is highly expressed both in whole blood and in brain tissues (particularly cerebellar hemisphere and cerebellum according to GTEx). While *CAMKK2* has not been implicated in a BP TWAS, the large PGC GWAS points toward calcium channel signaling as a potential therapeutic target for BP,^27^ and indeed a loss-of-function mutation in this gene has been previously linked to BP status.^49^ We consider these to be potential BP-relevant genes that are interesting candidates for experimental validation.

We replicated previous findings that immune cell type composition is impacted by lithium use rather than BP status. We also replicated several previously reported genes that are differentially expressed in whole blood in response to lithium, in addition to reporting novel lithium-response genes. Although lithium has been prescribed as a mood stabilizer for decades, its precise mechanism of action is still unclear.^50^ Lithium has been shown to increase the activity of the transcription factor CREB (cAMP response element-binding protein),^51^ a protein involved in neuronal plasticity.^52^ Here, we found that *ATF4*, an eGene in all cell types and the bulk, which encodes for CREB-2, has opposite directions of effect in T cell types than in the other immune cell types or bulk. We found a similar pattern for the *AKT1* (Rho-family-alpha serine/threonine-protein kinase) eGene. AKT1 protein levels in brain tissue have been previously associated with both schizophrenia and bipolar disorder, and although genetic associations exist,^53^ they do not pass genome-wide multiple testing correction.

While we find promising lines of evidence that immune cell type specific expression is useful for discovering candidate brain-relevant genes, there are several limitations to our study. Firstly, while our cohort had an ample number of BP patients, the number of SCZ samples was much lower, and thus underpowered for a diagnosis-specific analysis. Furthermore, we only test SNP-gene pairs in *cis*, whereas *trans* eQTLs are known to be more context-specific,^54^ so we miss distal associations that are potentially biologically relevant to the phenotypes of interest. By using computationally-derived expression estimates, there is a greater possibility for spurious associations that are not related to biology, dependent on the specific method of decomposition/deconvolution chosen. Also by using low-coverage RNA-seq, we may be missing important eGenes that are not as highly expressed in blood. Finally, our study consists of all European-ancestry individuals, but to gain a more comprehensive and inclusive understanding of the biology between immune cell types and psychiatric conditions, in addition to better fine-mapping these eQTL, many more samples of diverse ancestries need to be analyzed in future work.

Collectively, this suggests that while the bulk whole blood gene expression provides a greater number of significant findings overall, cell type specific expression allows us to observe additional biological mechanisms that are not possible to capture when only using gene expression measures from bulk alone.

## METHODS

### Cohort description

The samples included are from a study with individuals ascertained for bipolar disorder (BP) or Schizophrenia (SCZ). The cohort consists of 1,045 individuals with BP, 84 individuals with SCZ, and 601 controls with whole blood RNA-seq and corresponding genotypes (N=1,730 after excluding first degree relatives) included for all individuals.

### Bulk RNA-Sequencing

Bulk RNA-sequencing was performed at the UCLA Neurogenomics Core, using the TruSeq Stranded plus rRNA and GlobinZero library preparation method, as described previously.^10^ We used FASTQC to visually inspect the read quality from the lower-coverage whole blood RNA-Seq (5.9M reads/sample). We then used kallisto^55^ to pseudoalign reads to the GRCh37 gencode transcriptome (v33) and quantify estimates for transcript expression. We aggregated transcript counts to obtain gene level read counts using scripts from the GTEx consortium (https://github.com/broadinstitute/gtex-pipeline).

### Genotyping pipeline

Genotypes for the individuals included in the cohort were obtained from the following platforms: OmniExpressExome (N = 816), Psych Chip (N = 522), COEX (N = 162), Illumina550 (N=19), and Global Screening Array (N=211). Given that the SNP-genotype data came from numerous studies, the number of overlapping SNPs across all platforms was < 80k, prompting us to perform imputation separately for each genotyping platform, as previously described in Schwarz, et. al. 2022. Briefly, genotypes were first filtered for Hardy-Weinberg equilibrium p value < 1.0e-6 for controls and p value < 1.0e-10 for cases, with minor allele frequency (MAF) > 0.01, then were imputed using the 1000 Genomes Project phase 3 reference panel^56^ by chromosome using RICOPILI v.1^57^ separately per genotyping platform, then subsequently merged. Imputation quality was assessed by filtering variants where genotype probability > 0.8 and INFO score > 0.1. We restricted it to only autosomal chromosomes due to sex chromosome dosage, as commonly done.^58^

### Cell type proportion estimation

We estimated the proportion of cell types of the bulk whole blood RNA-seq datasets using CIBERSORTx, with batch correction applied and LM22 signature matrix as the reference gene expression profile. The LM22 signature matrix uses 547 genes to distinguish between 22 human hematopoietic cell phenotypes, though here we restrict to 8 cell types with proportions > 0.02.

Complete blood counts (CBC) lab tests from the clinic were provided for a subset of the cohort (N=143), providing us ground truth measures (in units of 10^9^ cells per liter) for neutrophils, lymphocytes, monocytes, basophils, and eosinophils. To make the counts comparable to the proportions outputted by CIBERSORTx, we divided the counts of the cell type of interest by the sum of counts across all cell types in an individual, providing the count ratio shown in Supplementary Figure 1.

### Cell type expression estimation

We log2-transformed the matrix of bulk TPM measures before inputting into bMIND since the largest expression measure was greater than 50 TPM. Using the cell type proportions derived from CIBERSORTx in conjunction with these log-transformed bulk expression measures, we used bMIND in order to derive cell type expression estimates, with flag np=TRUE.

### bMIND derived estimates and cis-eQTL mapping

Using output from bMIND, we transformed expression estimates from log2(TPM) to counts using sequencing library sizes, restricting to sufficiently expressed genes (estimated count > 1.0 in 40% of individuals). Expression estimates were then standardized (mean = 0) then performed cis-eQTL analysis mapping using QTLTools, using a defined window of 1 Mb both up and downstream of every gene’s TSS, for sufficiently expressed genes (TPM > 0.1 in 20% of individuals). We run the eQTL analysis in permutation pass mode (1000 permutations, and perform multiple testing corrections using the q value FDR procedure, correcting at 5% unless otherwise specified. We then restrict associations to the top (or leading) SNP per eGene.

### TWAS and colocalization

We used the FUSION pipeline to perform TWAS on the normalized cell type specific expression estimates and normalized bulk expression measures, residualizing each expression matrix by its first 50 principal components to account for variation due to technical (non-biological) factors. Imputed genotypes were restricted to those that overlap with the 1000 Genomes LD reference panel, providing 272,652 SNPs on which to perform the analysis. A window of 500kb upstream and 500kb downstream of the lead SNP for each eQTL was used as the cis-region to be tested. Gene-trait pairs were selected based on the best performing model after five-fold cross validation, including for Best Unbiased Linear Predictor (BLUP), elastic net (ENET), Least Absolute Shrinkage and Selection Operator (LASSO), and just using the top SNP. We tested for colocalization of GWAS and eQTLs using the –coloc flag within the FUSION/TWAS pipeline. Colocalization is only performed in those gene-trait associations with p < 0.05. In each cell type, we tested eGenes with a significant association between expression and SNP (**Tables 4 and 5**). We report SNPs with a colocalization probability (PP4) > 0.80.

### Cell type specific regressions using estimated cell type proportions and gene expression

We built logistic regression models to evaluate the effect of cell type proportion on case/control status, and lithium use status within only the BP cases. These models included the proportion of one cell type at a time, along with covariates including age, sex, RNA concentration, and RNA integrity number (RIN) as predictors. In testing the differences in cell type proportions between different binary outcomes, we used the glm() function in R with family=binomial.

We also used logistic regression to predict either case control status or lithium use (only in BP cases) from cell type expression estimates after residualizing for 50 expression PCs. Variable numbers of genes were included based on genes with most variance per cell type, using a range of 100 to 1000 genes with an interval of 100. Covariates include age, sex, RNA RIN, RNA concentration, and cell type proportion estimates. A random 70% of individuals were sampled to use for training, and 30% for testing the prediction.

### Electronic Medical Record Validation Cohort

ATLAS is an opt-in biobank that enrolls patients when they visit UCLA for a blood draw. ATLAS is a diverse biobank that includes patients from a variety of genetic ancestries that live across the greater Los Angeles region.^59^ Registered ATLAS researchers can access deidentified electronic health record data for patients, consisting of outpatient and inpatient encounters, including information on diagnoses, procedure orders, laboratory orders, and prescription orders. As of 2022, there were approximately 50,000 participants enrolled in ATLAS. A complete description of the ATLAS project and data is available in ^44^.

Bipolar patients were identified in ATLAS using the diagnosis table. The bipolar phenotype was defined as any patient who had at least one diagnosis of any of the ICD 10 codes included in the bipolar Phecode Map 1.2.^60^ Neutrophil counts (measured as 10^3^ counts/µL) were determined using test results for complete blood count laboratory orders. We restricted this analysis to those individuals with self-reported European ancestry. To prevent severe outliers from biasing results, test results with a neutrophil count greater than 2 standard deviations from the median count value in all bipolar patients were removed. Lithium prescription orders were found by querying the prescription order table for medications of any dose or format that were classified as psychiatric medication and had the generic name lithium.

Neutrophil count data for patients with a bipolar Phecode were separated into three categories: tests administered before the patient was prescribed lithium, tests administered after the first lithium prescription order, and tests for patients without a lithium prescription order. Since many patients had multiple complete blood count orders, the median neutrophil count per patient per category was calculated. Median neutrophil counts were compared between bipolar patients after their first lithium prescription and bipolar patients without a lithium prescription using a logistic regression (implemented in R). Max age and sex were used as covariates. For the subset of patients who had complete blood count tests taken before and after a lithium prescription order, we used a paired Wilcoxon rank test to increase power, implemented in R using the wilcox.test(paired=TRUE) command.

### Interaction model

To test whether there exists an interaction between SNP-lithium usage, we included an interaction component in the regression model, as such: *y* = β* *X* + β * ℓ + β * (*X* + ℓ) + *covariates* where *X* refers to the genotype at a particular SNP, and *l* refers to lithium use.

### Differential expression analysis

We used the limma eBayes function with trend=true to conduct differential expression tests in the bulk dataset. We include only those genes with at least 1 TPM in at least 436 individuals (about 25% of the total 1,730 individuals included in the analysis), leaving 17,194 genes to be tested. We then log2-transform this matrix and compute the first 50 expression principal components to be included as covariates. In the lithium user vs non-user analysis, only cases were included to avoid confounding effects caused by disease status, while in the case-control analysis, all individuals diagnosed with BP or SCZ were included as cases and non-affected individuals included as controls.

For the cell-type specific differential expression analysis, we use the bmind_de() function as included in the bMIND software package. To keep the methods comparable to the bulk analysis, we also use the log2-transformed expression measures as input along with the first 50 expression PCs as covariates.

## Data Availability

The lower-coverage RNA-seq and the corresponding genotypes generated and analyzed during this study have been deposited in dbGAP (accession number phs002856.v1).

## Supporting information

Differential Expression Supplementary Tables

TWAS Supplemental Tables

## Acknowledgements

We are greatly appreciative to those individuals who donated the blood samples on which this study was based. We gratefully acknowledge the Institute for Precision Health, participating patients from the UCLA ATLAS Precision Health Biobank, UCLA David Geffen School of Medicine, UCLA Clinical and Translational Science Institute, and UCLA Health. T.B. was supported by the NIH (grant no. 5T32HG002536-19). We also would like to thank Tanner Waters for his meaningful discussions on this work. This research was supported by the National Institute of Mental Health of the National Institutes of Health under award no. 2R01MH115676-06, former number 5R01MH115676-05. The content is solely the responsibility of the authors and does not necessarily represent the official views of the National Institutes of Health.

## Supplementary Figures

**Supplementary Figure 1:**
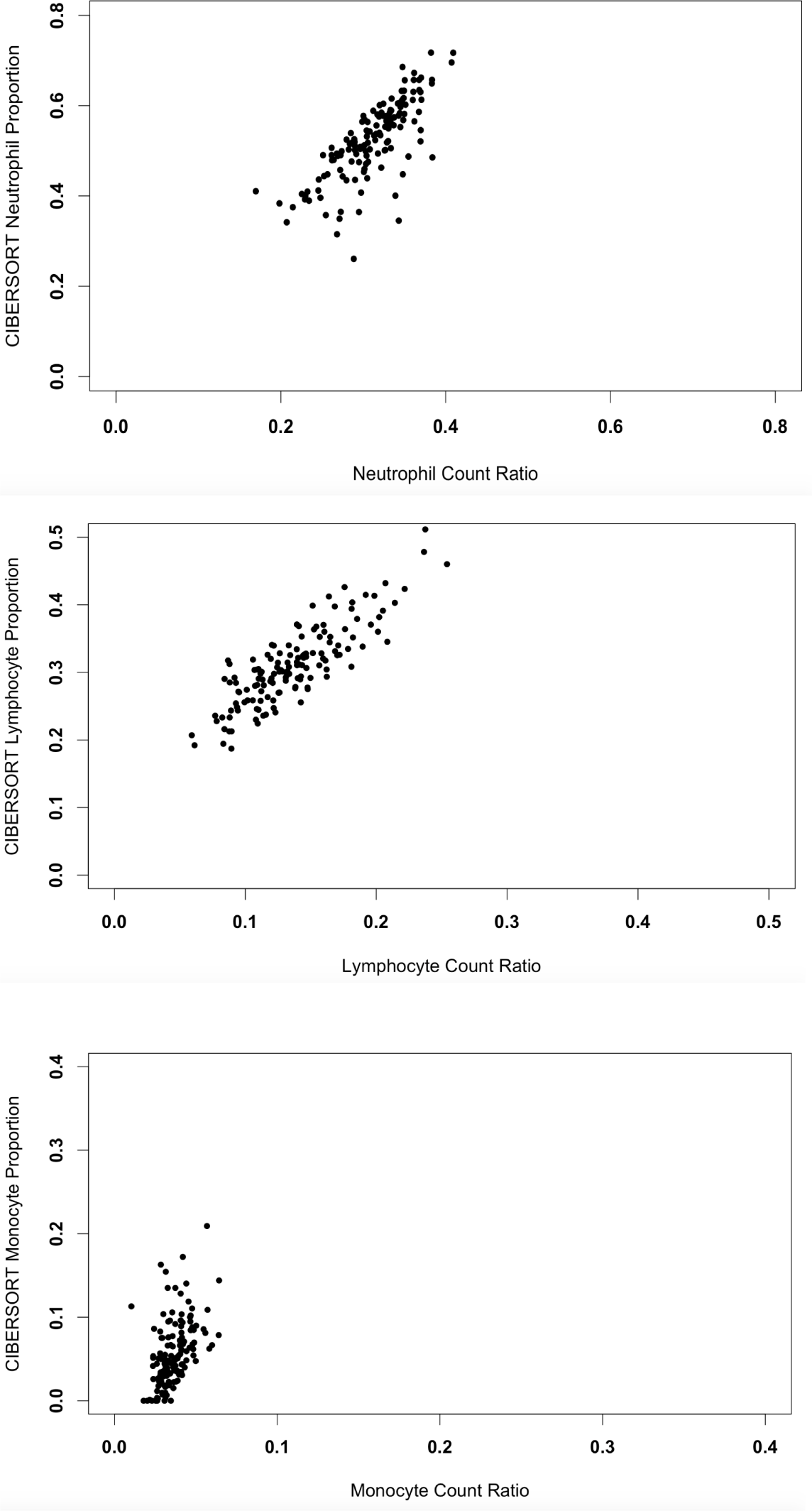
Scatterplots of CIBERSORTx-estimated cell type proportions vs complete blood count proportions. We find generally high concordance between computationally estimated and measured ground truth cell type proportions using a subset of our cohort. Pearson’s correlation R2 for neutrophils = 0.76, for lymphocytes = 0.85, for monocytes = 0.48.

**Supplementary Figure 2:**
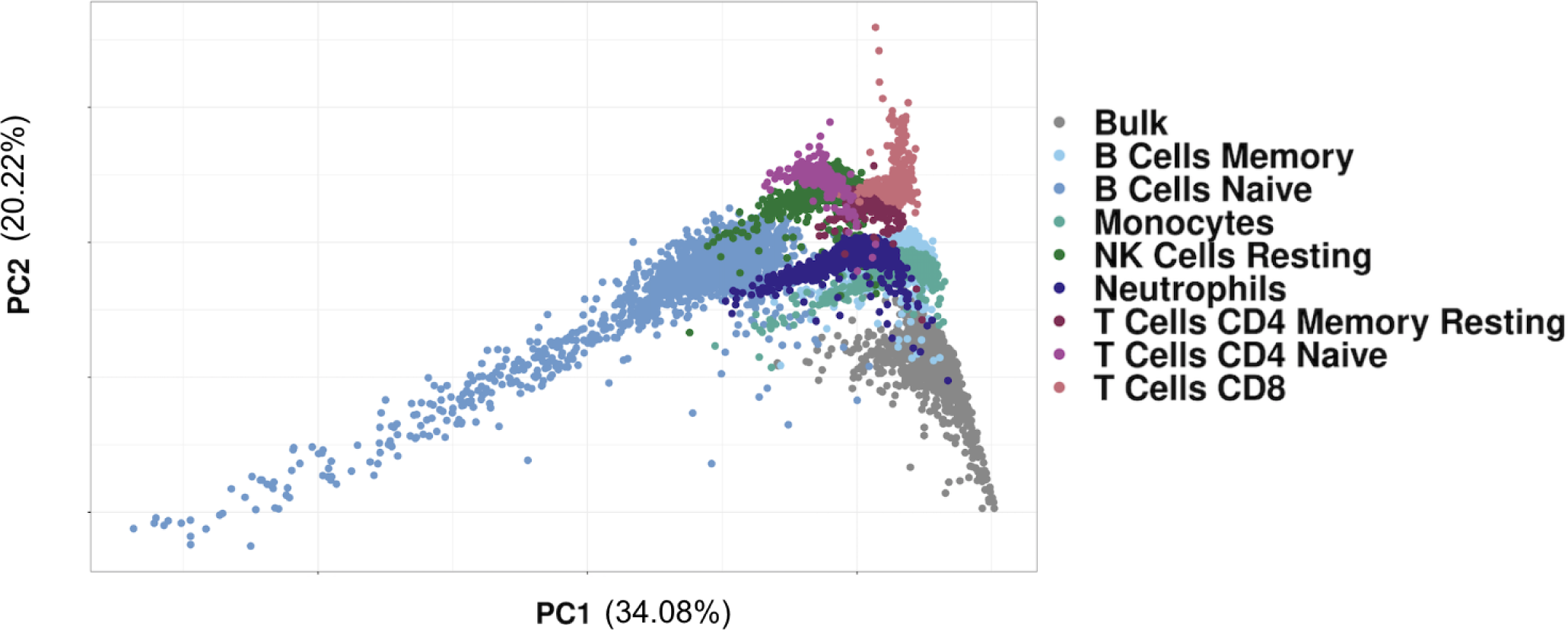
PCA of cell type expression.

**Supplementary Figure 3:**
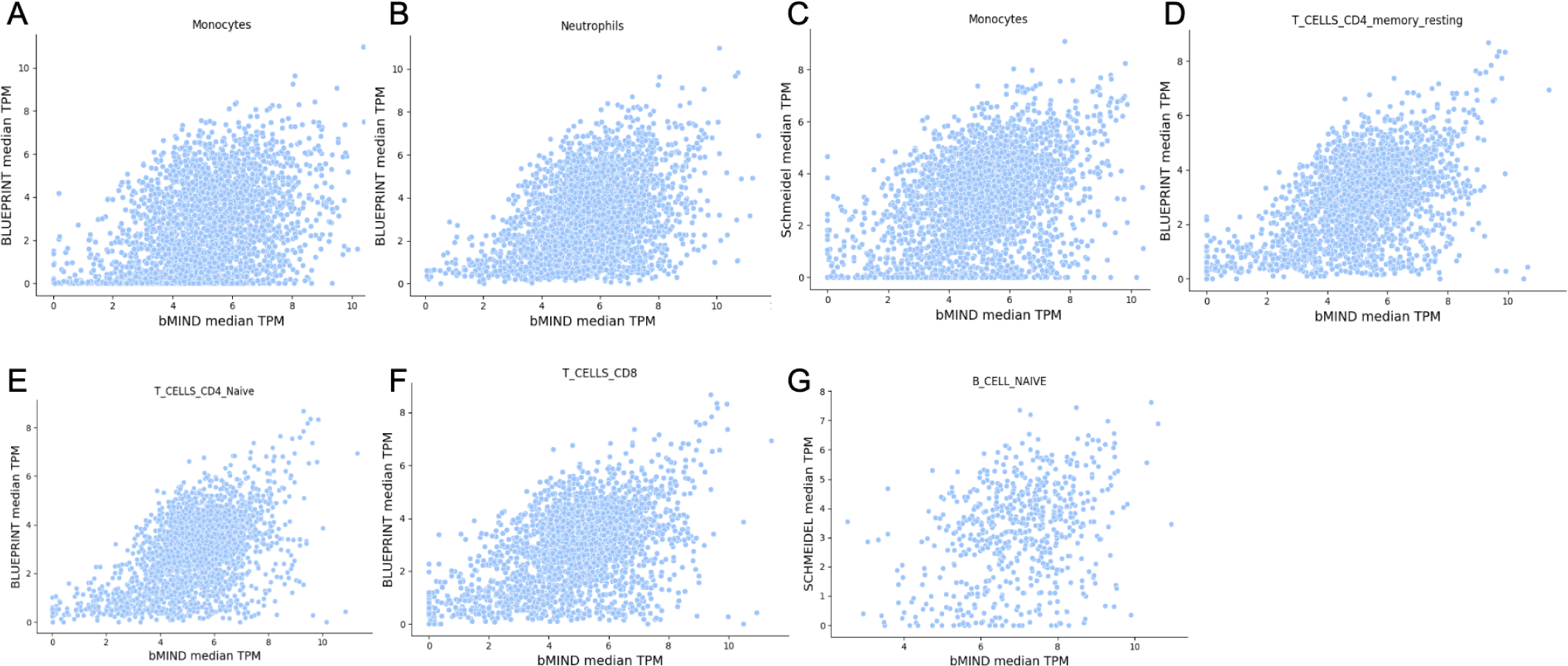
Scatterplots of expression estimated from bulk vs single-cell. Using two scRNA-Seq datasets as references, we compare the median TPM values for genes detected as eQTLs using both scRNA-Seq and computationally deconvoluted bulk RNA-Seq.

**Supplementary Figure 4:**
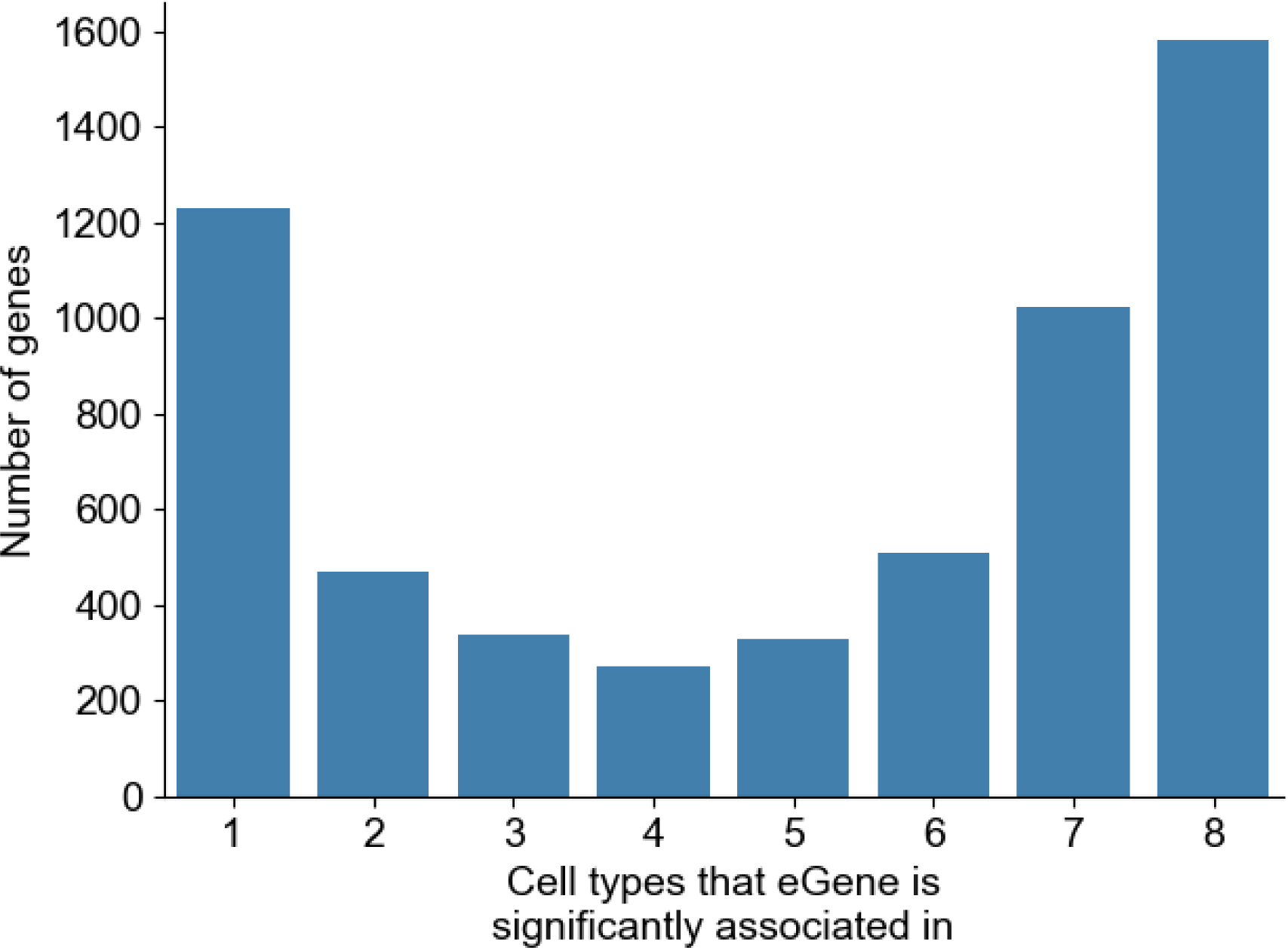
Distribution of shared eGenes across cell type contexts.

**Supplementary Figure 5:**
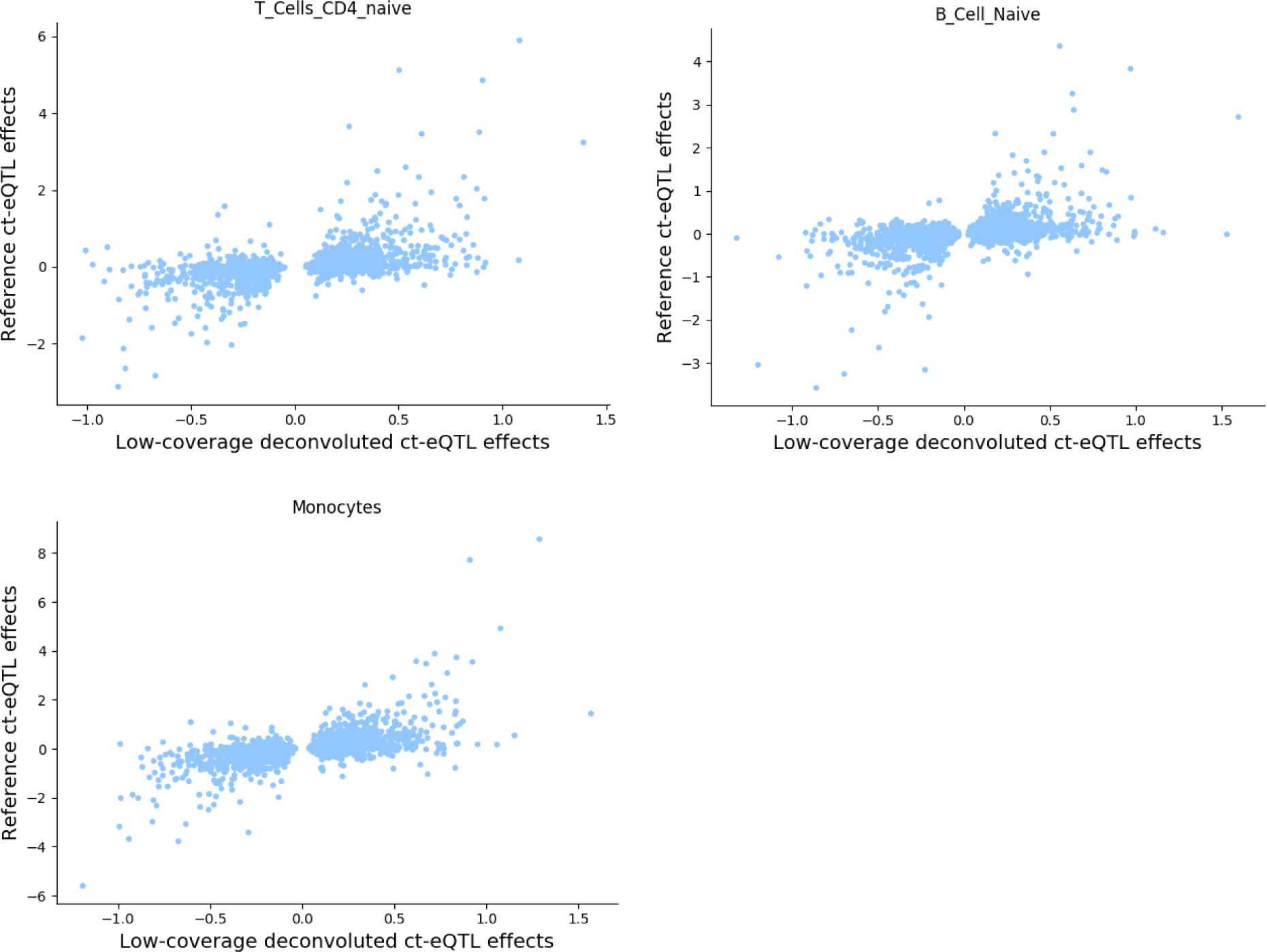
Effect size correlations between reference single cell eQTL and the deconvoluted eQTL. T cells CD4 naive R2 = 0.27; B cells naive R2 = 0.22; Monocytes R2 = 0.36.

**Supplementary Figure 6:**
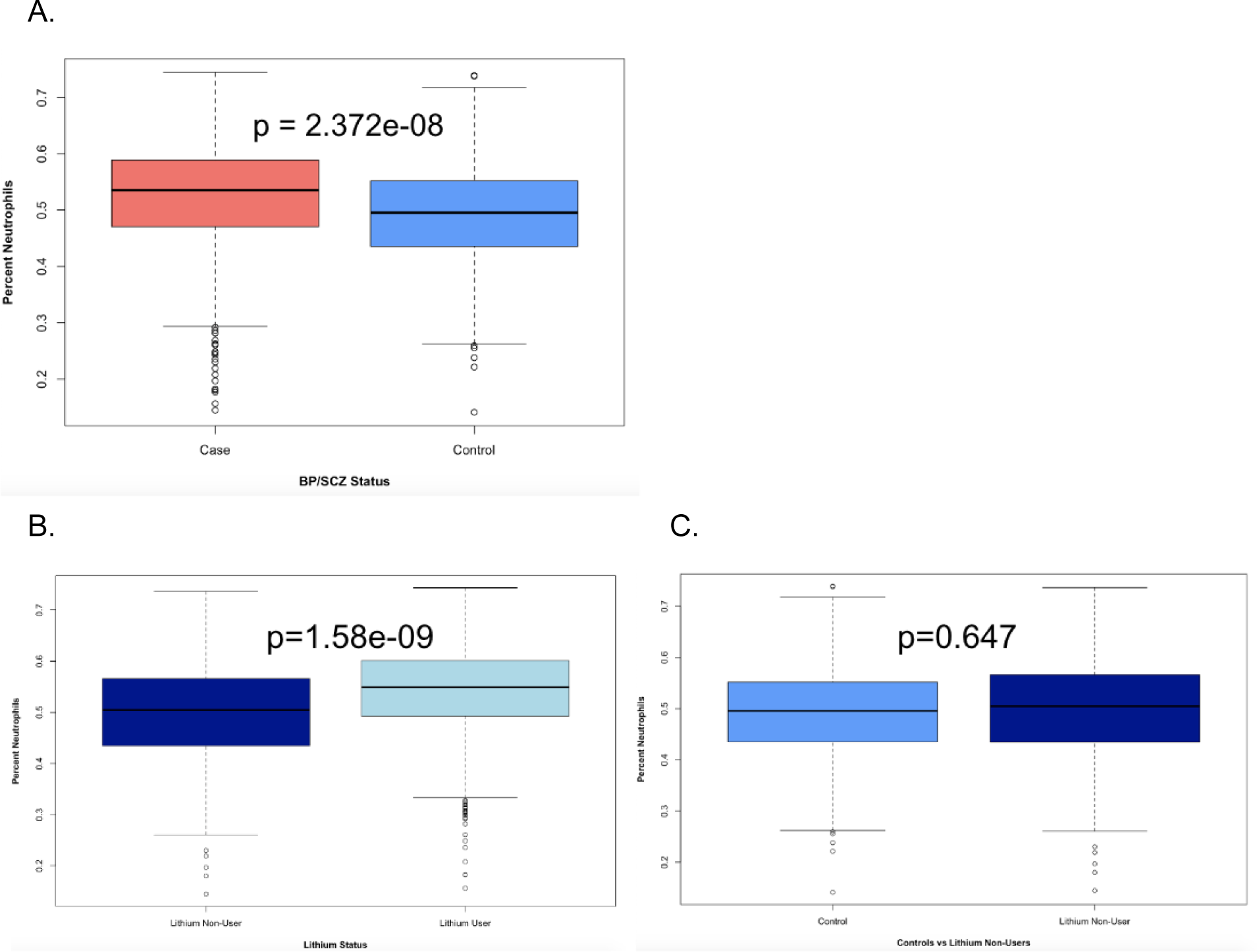
Neutrophil count elevated for lithium users. A. Difference in neutrophil proportion (after accounting for covariates including age, sex, RIN, and RNA concentration) between BP/SCZ cases and controls. B. Difference in neutrophil proportion between lithium users and non-users (after accounting for covariates), only within BP cases. C. Difference in neutrophil proportion between lithium non-users and controls (after accounting for covariates).

**Supplementary Figure 7:**
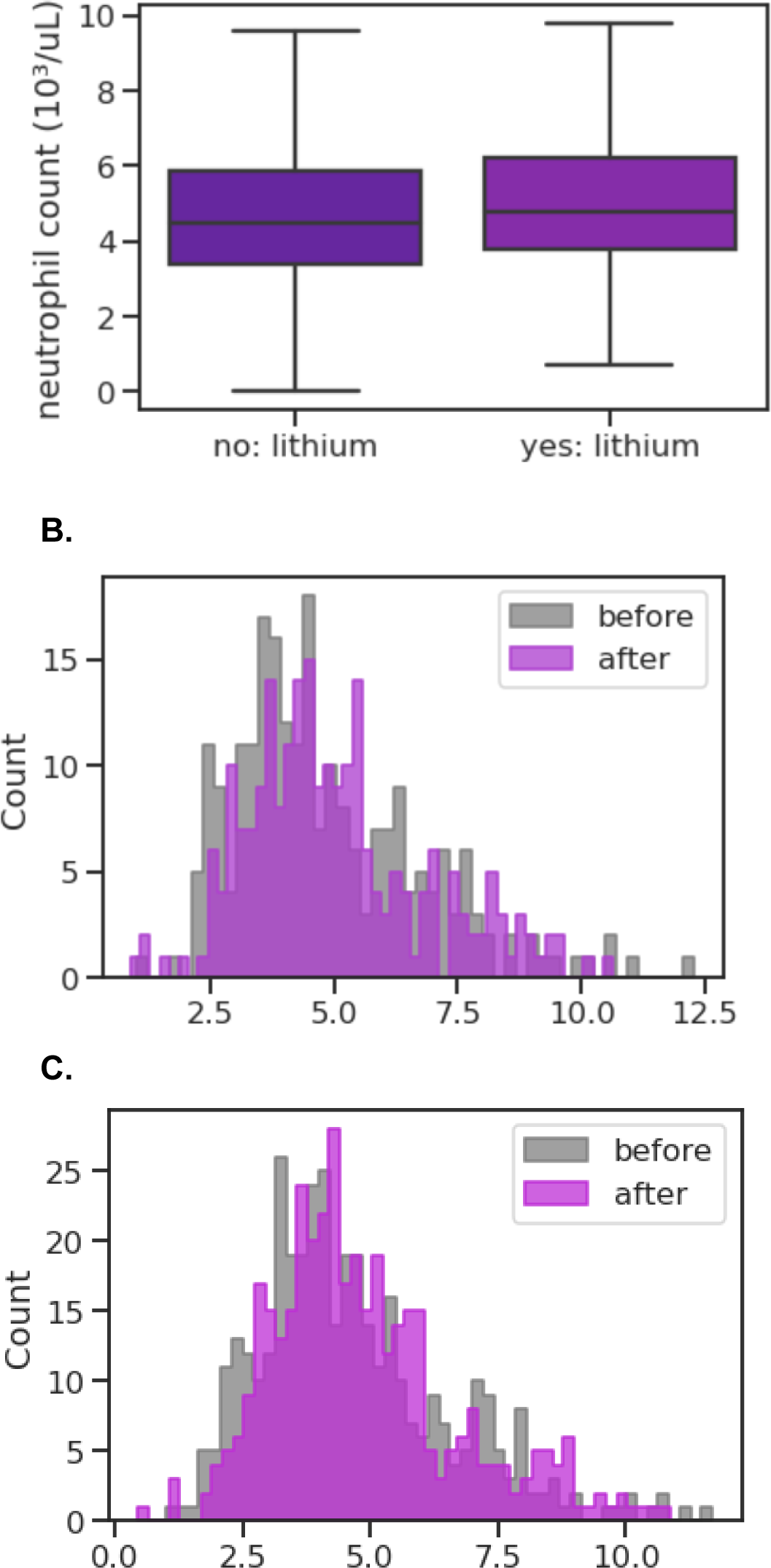
Neutrophil count elevated for lithium users in UCLA ATLAS. A. Median neutrophil count across self-reported European patients, with covariate correction for age and sex (p=2.09e-7). B. Neutrophil count distribution across self-reported European patients (n=X) before and after lithium prescription (Wilcoxon p=X). C. Neutrophil count distribution across patients (n=382, all ancestries) before and after lithium prescription (Wilcoxon p=0.0228).

**Supplementary Figure 8:**
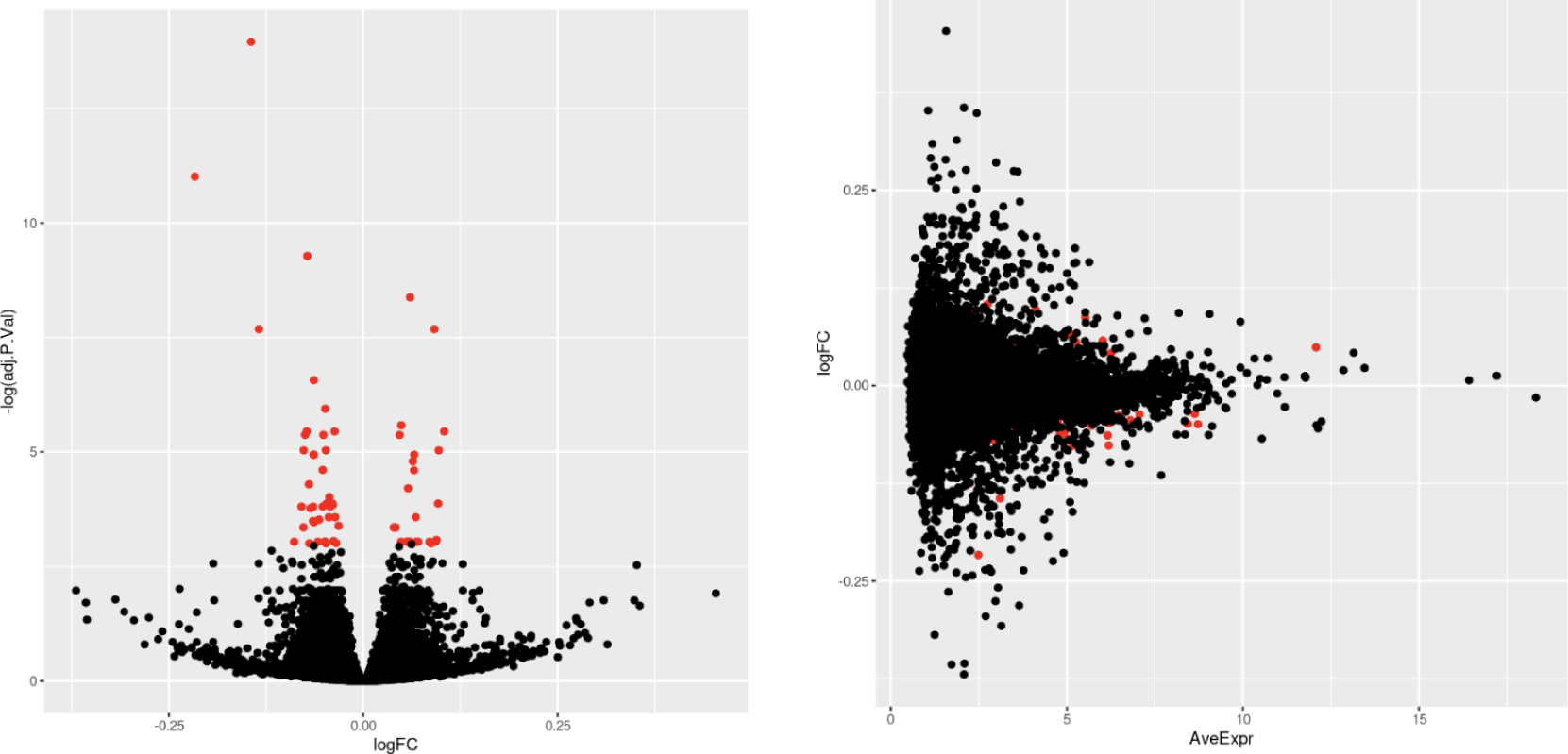
Differential gene expression results for BP or SCZ cases vs controls. (left) Volcano plot which highlights differentially expressed genes (FDR < 0.05) in red (N=64 total differentially expressed genes). (right) Average expression of each gene vs the log fold change (logFC) of each gene, with differentially expressed genes highlighted in red.

**Supplementary Table 1:** Correlations of median expression between reference single cell RNA-Seq datasets and computationally derived expression estimates. Restricting to the genes identified as eGenes using both the single cell RNA-Seq reference dataset and the computationally derived cell type expression, we report R2 values for the median TPM for genes across samples.

| Cell type | Reference | $R^2$ | Number of genes |
| --- | --- | --- | --- |
| Monocytes | BLUEPRINT | 0.14 | 2896 |
| Neutrophils | BLUEPRINT | 0.16 | 3239 |
| CD4 Memory T Cells | BLUEPRINT | 0.26 | 2504 |
| CD4 Naive T Cells | BLUEPRINT | 0.27 | 2504 |
| CD8 T Cells | BLUEPRINT | 0.24 | 2504 |
| B Cell Naive | Schmeidel | 0.11 | 624 |
| Monocytes | Schmeidel | 0.15 | 2896 |
| Monocytes* | Schmeidel/BLUEPRINT | 0.22 | 2836 |

**Supplementary Table 2:** Linear models for lithium usage prediction.

| Cell type | Model $R^2$ | Effect size | SE | p-value |
| --- | --- | --- | --- | --- |
| Monocytes | 0.06 | -1.28 | 0.36 | <0.001 |
| Neutrophils | 0.09 | 1.08 | 0.15 | <0.001 |
| CD4 Memory T Cells | 0.06 | -1.68 | 0.42 | <0.001 |
| CD4 Naive T Cells | 0.06 | -1.52 | 0.36 | <0.001 |
| CD8 T Cells | 0.06 | -2.09 | 0.55 | <0.001 |
| B Cell Naive | 0.06 | -3.06 | 0.79 | <0.001 |
| B Cell Memory | 0.05 | 0.59 | 1.09 | 0.59 |
| Resting NK Cells | 0.06 | -1.92 | 0.52 | <0.001 |

**Supplementary Table 3:** Number of eGenes differentially regulated by Lithium. Using an FDR cut-off of p < 0.10, we look at the number of eGenes with a significant SNP-lithium interaction. “Same-direction” Li-eGenes have the same direction of effect sizes between lithium users and nonusers, and “opposite-direction” Li-eGenes have the opposite direction of effect sizes between lithium users and nonusers.

| Cell type | Number of Li-eGenes | Number of “same-direction” Li-eGenes | Number of “opposite-direction” Li-eGenes |
| --- | --- | --- | --- |
| Naive B cells | 24 | 4 | 20 |
| Memory B cells | 15 | 3 | 12 |
| CD8 T cells | 2 | 1 | 1 |
| Naive CD4 T cells | 25 | 1 | 24 |
| Memory CD4 T cells | 2 | 0 | 2 |
| Resting NK cells | 5 | 0 | 5 |
| Monocytes | 34 | 3 | 31 |
| Neutrophils | 3 | 1 | 2 |

## References

1. Zeng, B. et al. Multi-ancestry eQTL meta-analysis of human brain identifies candidate causal variants for brain-related traits. Nat. Genet. 54, 161–169 (2022).

2. Zhang, J. & Zhao, H. eQTL Studies: from Bulk Tissues to Single Cells. ArXiv (2023).

3. Finucane, H. K. et al. Partitioning heritability by functional annotation using genome-wide association summary statistics. Nat. Genet. 47, 1228–1235 (2015).

4. Kim-Hellmuth, S. et al. Cell type-specific genetic regulation of gene expression across human tissues. Science 369, (2020).

5. Mandric, I. et al. Optimized design of single-cell RNA sequencing experiments for cell-type-specific eQTL analysis. Nat. Commun. 11, 5504 (2020).

6. Cobos, F. A., Alquicira-Hernandez, J., Powell, J. E., Mestdagh, P. & De Preter, K. Benchmarking of cell type deconvolution pipelines for transcriptomics data. Nature Communications vol. 11 Preprint at https://doi.org/10.1038/s41467-020-19015-1 (2020).

7. Jin, H. & Liu, Z. A benchmark for RNA-seq deconvolution analysis under dynamic testing environments. Genome Biol. 22, 102 (2021).

8. Newman, A. M. et al. Determining cell type abundance and expression from bulk tissues with digital cytometry. Nat. Biotechnol. 37, 773–782 (2019).

9. Wang, J., Roeder, K. & Devlin, B. Bayesian estimation of cell type-specific gene expression with prior derived from single-cell data. Genome Res. (2021) doi:10.1101/gr.268722.120.

10. Schwarz, T. et al. Powerful eQTL mapping through low coverage RNA sequencing. Preprint at https://doi.org/10.1101/2021.08.08.455466.

11. Khandaker, G. M., Dantzer, R. & Jones, P. B. Immunopsychiatry: important facts. Psychol. Med. 47, 2229–2237 (2017).

12. Qi, T. et al. Identifying gene targets for brain-related traits using transcriptomic and methylomic data from blood. Nat. Commun. 9, 2282 (2018).

13. Werner, M. C. F. et al. Immune marker levels in severe mental disorders: associations with polygenic risk scores of related mental phenotypes and psoriasis. Transl. Psychiatry 12, 1–8 (2022).

14. Psychiatric genome-wide association study analyses implicate neuronal, immune and histone pathways. Nat. Neurosci. 18, 199–209 (2015).

15. Le Clerc, S. et al. HLA-DRB1 and HLA-DQB1 genetic diversity modulates response to lithium in bipolar affective disorders. Sci. Rep. 11, 17823 (2021).

16. Chernecky, C. C. & Berger, B. J. Laboratory Tests and Diagnostic Procedures. (W.B. Saunders Company, 1997).

17. George-Gay, B. & Parker, K. Understanding the complete blood count with differential. J. Perianesth. Nurs. 18, 96–114; quiz 115–7 (2003).

18. Schmiedel, B. J. et al. Single-cell eQTL analysis of activated T cell subsets reveals activation and cell type-dependent effects of disease-risk variants. Science immunology 7, (2022).

19. Chen, L. et al. Genetic Drivers of Epigenetic and Transcriptional Variation in Human Immune Cells. Cell 167, 1398–1414.e24 (2016).

20. Moffat, J. J., Ka, M., Jung, E.-M., Smith, A. L. & Kim, W.-Y. The role of MACF1 in nervous system development and maintenance. Semin. Cell Dev. Biol. 69, 9–17 (2017).

21. Bryois, J. et al. Cell-type specific cis-eQTLs in eight brain cell-types identifies novel risk genes for human brain disorders. Preprint at https://doi.org/10.1101/2021.10.09.21264604.

22. Westra, H.-J. et al. Cell Specific eQTL Analysis without Sorting Cells. PLoS Genet. 11, e1005223 (2015).

23. Schmiedel, B. J. et al. Impact of Genetic Polymorphisms on Human Immune Cell Gene Expression. Cell vol. 175 1701–1715.e16 Preprint at https://doi.org/10.1016/j.cell.2018.10.022 (2018).

24. Kerimov, N. et al. A compendium of uniformly processed human gene expression and splicing quantitative trait loci. Nat. Genet. 53, 1290–1299 (2021).

25. Gusev, A. et al. Integrative approaches for large-scale transcriptome-wide association studies. Preprint at https://doi.org/10.1101/024083.

26. Giambartolomei, C. et al. Bayesian test for colocalisation between pairs of genetic association studies using summary statistics. PLoS Genet. 10, e1004383 (2014).

27. Mullins, N. et al. Genome-wide association study of more than 40,000 bipolar disorder cases provides new insights into the underlying biology. Nat. Genet. 53, 817–829 (2021).

28. Consortium, T. S. W. G. of T. P. G., The Schizophrenia Working Group of the Psychiatric Genomics Consortium, Ripke, S., Walters, J. T. R. & O’Donovan, M. C. Mapping genomic loci prioritises genes and implicates synaptic biology in schizophrenia. Preprint at https://doi.org/10.1101/2020.09.12.20192922.

29. Wray, N. R. et al. Genome-wide association analyses identify 44 risk variants and refine the genetic architecture of major depression. Nat. Genet. 50, 668–681 (2018).

30. Walters, R. K. et al. Transancestral GWAS of alcohol dependence reveals common genetic underpinnings with psychiatric disorders. Nat. Neurosci. 21, 1656–1669 (2018).

31. Johnson, E. C. et al. A large-scale genome-wide association study meta-analysis of cannabis use disorder. Lancet Psychiatry 7, 1032–1045 (2020).

32. Watanabe, K. et al. Author Correction: A global overview of pleiotropy and genetic architecture in complex traits. Nat. Genet. 52, 353 (2020).

33. Jansen, P. R. et al. Genome-wide analysis of insomnia in 1,331,010 individuals identifies new risk loci and functional pathways. Nat. Genet. 51, 394–403 (2019).

34. Demontis, D. et al. Discovery of the first genome-wide significant risk loci for attention deficit/hyperactivity disorder. Nat. Genet. 51, 63–75 (2019).

35. Jansen, I. E. et al. Genome-wide meta-analysis identifies new loci and functional pathways influencing Alzheimer’s disease risk. Nat. Genet. 51, 404–413 (2019).

36. Akbarian, S. et al. The PsychENCODE project. Nature Neuroscience vol. 18 1707–1712 Preprint at https://doi.org/10.1038/nn.4156 (2015).

37. Bentham, J. et al. Genetic association analyses implicate aberrant regulation of innate and adaptive immunity genes in the pathogenesis of systemic lupus erythematosus. Nat. Genet. 47, 1457–1464 (2015).

38. van der Harst, P. et al. Seventy-five genetic loci influencing the human red blood cell. Nature 492, 369–375 (2012).

39. Bycroft, C. et al. The UK Biobank resource with deep phenotyping and genomic data. Nature 562, 203–209 (2018).

40. Wang, Y.-F. et al. Identification of 38 novel loci for systemic lupus erythematosus and genetic heterogeneity between ancestral groups. Nat. Commun. 12, 772 (2021).

41. Eames, H. L., Corbin, A. L. & Udalova, I. A. Interferon regulatory factor 5 in human autoimmunity and murine models of autoimmune disease. Transl. Res. 167, 167–182 (2016).

42. Akkouh, I. A. et al. Exploring lithium’s transcriptional mechanisms of action in bipolar disorder: a multi-step study. Neuropsychopharmacology vol. 45 947–955 Preprint at https://doi.org/10.1038/s41386-019-0556-8 (2020).

43. Krebs, C. E. et al. Whole blood transcriptome analysis in bipolar disorder reveals strong lithium effect. Psychol. Med. 50, 2575–2586 (2020).

44. Johnson, R. et al. Leveraging genomic diversity for discovery in an electronic health record linked biobank: the UCLA ATLAS Community Health Initiative. Genome Medicine vol. 14 Preprint at https://doi.org/10.1186/s13073-022-01106-x (2022).

45. Zhou, W. et al. Global Biobank Meta-analysis Initiative: Powering genetic discovery across human disease. Cell Genom 2, 100192 (2022).

46. Smyth, G. K. limma: Linear Models for Microarray Data. Bioinformatics and Computational Biology Solutions Using R and Bioconductor 397–420 Preprint at https://doi.org/10.1007/0-387-29362-0_23.

47. Munkholm, K., Peijs, L., Vinberg, M. & Kessing, L. V. A composite peripheral blood gene expression measure as a potential diagnostic biomarker in bipolar disorder. Transl. Psychiatry 5, e614 (2015).

48. Lisé, M.-F. et al. Myosin-Va-interacting protein, RILPL2, controls cell shape and neuronal morphogenesis via Rac signaling. J. Cell Sci. 122, 3810–3821 (2009).

49. Ling, N. X. Y. et al. Functional analysis of an R311C variant of Ca ^2^ -calmodulin-dependent protein kinase kinase-2 (CaMKK2) found as a de novo mutation in a patient with bipolar disorder. Bipolar Disorders vol. 22 841–848 Preprint at https://doi.org/10.1111/bdi.12901 (2020).

50. Alda, M. LITHIUM IN THE TREATMENT OF BIPOLAR DISORDER: PHARMACOLOGY AND PHARMACOGENETICS. Mol. Psychiatry 20, 661 (2015).

51. Grimes, C. A. & Jope, R. S. CREB DNA binding activity is inhibited by glycogen synthase kinase-3 beta and facilitated by lithium. J. Neurochem. 78, (2001).

52. Sakamoto, K., Karelina, K. & Obrietan, K. CREB: a multifaceted regulator of neuronal plasticity and protection. J. Neurochem. 116, 1–9 (2011).

53. Karege, F. et al. Association of AKT1 gene variants and protein expression in both schizophrenia and bipolar disorder. Genes Brain Behav. 9, 503–511 (2010).

54. GTEx Consortium et al. Genetic effects on gene expression across human tissues. Nature 550, 204–213 (2017).

55. Bray, N. L., Pimentel, H., Melsted, P. & Pachter, L. Near-optimal probabilistic RNA-seq quantification. Nat. Biotechnol. 34, 525–527 (2016).

56. 1000 Genomes Project Consortium et al. A global reference for human genetic variation. Nature 526, 68–74 (2015).

57. Lam, M. et al. RICOPILI: Rapid Imputation for COnsortias PIpeLIne. Bioinformatics 36, 930–933 (2020).

58. Consortium, T. G. & The GTEx Consortium. The GTEx Consortium atlas of genetic regulatory effects across human tissues. Science vol. 369 1318–1330 Preprint at https://doi.org/10.1126/science.aaz1776 (2020).

59. Caggiano, C. et al. Health care utilization of fine-scale identity by descent clusters in a Los Angeles biobank. Preprint at https://doi.org/10.1101/2022.07.12.22277520.

60. Wu, P. et al. Mapping ICD-10 and ICD-10-CM Codes to Phecodes: Workflow Development and Initial Evaluation. JMIR Med Inform 7, e14325 (2019).

61. Farioli-Vecchioli, S. et al. Btg1 is Required to Maintain the Pool of Stem and Progenitor Cells of the Dentate Gyrus and Subventricular Zone. Front. Neurosci. 6, 124 (2012).

